# Neural posterior estimation for population genetics

**DOI:** 10.64898/2025.12.01.691638

**Authors:** Jiseon Min, Yuxin Ning, Nathaniel S. Pope, Franz Baumdicker, Andrew D. Kern

## Abstract

Simulation-based inference methods are increasingly being used in population genetics due to their flexibility and ability to be applied in settings where likelihood-based methods are intractable. Perhaps the best known such method is Approximate Bayesian Computation (ABC); however, its popularity is offset by its shortcomings which include computational expense and an unfortunate inability to *efficiently* fit models to high-dimensional summaries of the data. An alternative approach that solves these issues is supervised machine learning (ML); however, ML methods generally do not yield Bayesian uncertainty estimates of the quantities they predict. Here, we apply a recently introduced method, neural posterior estimation (NPE), that combines the best facets of ABC and supervised ML by training a neural network to estimate the posterior distribution of a population genetics model. We first compare neural posterior estimation with other inference methods for a variety of population genetic tasks, and show that neural posterior estimators yield posterior distributions with high accuracy and efficiency. We compare learned posterior distributions given raw genotypes and various summary statistics as input data. Additionally, we apply neural posterior estimation for demographic inference for simple and more complex models to highlight its application, including an analysis of demographic history in *Drosophila melanogaster*. Finally, we provide a user friendly workflow that enables others to perform neural posterior estimation on their own genetic data.

## Introduction

Population genetic inference typically proceeds via statistical estimation using a few well-described models (i.e., the Wright-Fisher model or its genealogical dual, the coalescent). Generally speaking, the key step is to describe how some summary of the data (e.g., the site frequency spectrum) depends on the parameters of a given evolutionary model. A familiar example is the software package `∂a∂i` (Gutenkunst et al., 2009), which uses a diffusion approximation to the Wright-Fisher model to estimate the population parameters that best fit an observed site frequency spectrum. The fitting occurs through maximum likelihood estimation, wherein the probability of the data (in this case, the site frequency spectrum) is optimized with respect to the parameters. Maximum likelihood estimation is a fundamental approach to statistical inference which allows uncertainty quantification via Fisher’s information matrix (or its generalization, the Godambe information matrix (Godambe, 1960; Varin et al., 2011))—the variance-covariance matrix that describes how much information the data holds for each parameter of the model— through the assumption of asymptotic normality of the estimators. Likelihood estimation has a long history in population genetics, and accordingly, there are many methods that use this approach, including for demographic inference (Adams and Hudson, 2004; Gutenkunst et al., 2009; Jouganous et al., 2017; Ragsdale and Gravel, 2019), estimation of recombination rates (Hudson, 2001; McVean et al., 2002; Chan et al., 2012; Spence and Song, 2019), and estimation of selection parameters (Bustamante et al., 2001, 2003; Nielsen et al., 2005) among other uses. While likelihood-based methods can be accurate, they are also limiting—model complexity and/or realism is often sacrificed when we depend on analytical results for inference.

An alternative approach relies on simulation to estimate the probability of observing certain features of a population genetic dataset and then to perform inference accordingly. Perhaps the best-known such approach in population genetics is Approximate Bayesian Computation (ABC; Beaumont et al. (2002)). Using ABC, a dataset is summarized by an informative set of features (e.g., heterozygosity, the number of haplotypes, etc.) and simulations of a given evolutionary scenario are performed using prior distributions of the parameters of interest. Each simulation is compared to the observed dataset, and either a rejection step or a regression step (or both) is used to estimate quantities from the posterior distribution of the parameters. ABC at this point has been well-studied and widely-applied (e.g., Thornton and Andolfatto, 2006; Laurent et al., 2011; Smith and Flaxman, 2020, among others); however, its popularity is offset by its shortcomings which include computational expense and an unfortunate inability to fit models to high-dimensional summaries of the data (Prangle, 2015). Several approaches have been proposed to mitigate this within the ABC framework, including semi-automatic summary statistic construction (Fearnhead and Prangle, 2012), partial least-squares regression (Wegmann et al., 2009), and subset selection (Joyce and Marjoram, 2008). At the same time, ABC highlights two substantial advantages of working with carefully chosen summary statistics in population genetics: they can be made interpretable in terms of classical theory and under some conditions they provide a principled, model-based route to inference with formal asymptotic guarantees. In what follows, we will use ABC primarily as a conceptual reference point for simulation-based Bayesian inference, rather than as a direct competitor that we benchmark against in detail.

Recently the field of population genetics has moved towards supervised ML and deep learning approaches which have been able to overcome many of the potential drawbacks of ABC (reviewed in Schrider and Kern, 2018; Huang et al., 2024). Through leveraging classical ML approaches, such as random forests and boosting, the associated “curse of dimensionality” with ABC feature selection can be side-stepped, allowing for a large number of potentially correlated summaries to be used without issue. Some early applications of this approach include methods for selective and demographic inference (Lin et al., 2011; Ronen et al., 2013; Schrider and Kern, 2015, 2016; Schrider et al., 2018), as well as for recombination rate estimation (Lin et al., 2013). These ML-based approaches have the additional benefit of amortization—once a model is sufficiently trained, no further computational investment is needed for doing estimation again and again on new data points. While that is so, classic ML methods don’t save the developer from the hard work of feature engineering—just as in ABC, one needs to proffer an appropriately informative set of summaries to the method for learning to occur.

To solve this problem, increasingly, population genetics has used so-called “end-to-end” learning as implemented by modern approaches to deep learning (DL). By using artificial neural network (ANN) architectures capable of automated feature extraction (Chan et al., 2018; Flagel et al., 2019; Adrion et al., 2020a), such as convolutional and recurrent neural networks (CNNs and RNNs respectively; reviewed in Huang et al. (2024)), a feature set needs not be hand-crafted, but instead *learned* by the method itself. While less interpretable than summary statistic based methods, this approach allows ANNs to automatically extract informative features from the data without human supervision. Indeed, this effort has been quite successful in population genetics over the last five or so years, and its applications have blossomed (e.g. Gower et al. (2021); Sanchez et al. (2021); Torada et al. (2019); López-Cortés et al. (2020)).

However, ABC outshines current deep learning approaches in population genetics in one crucial respect: an explicit quantification of the uncertainty surrounding parameter estimates in the form of a posterior distribution. Recent progress in training deep neural networks to produce posterior densities rather than point estimates promises to fuse the efficiency of DL for simulation-based inference with the principled probabilistic logic of Bayesian statistics (Papamakarios et al., 2019; Greenberg et al., 2019; Hermans et al., 2020). These neural posterior estimation (NPE) methods combine deep neural networks with flexible density estimators (e.g., normalizing flows) to learn a function that maps an input (such as a genotype array or a vector of summary statistics) directly to an approximation of the posterior distribution. Conceptually, NPE can be viewed as an amortized, learned analogue of the regression-adjustment step in ABC (where a regression model is fit to the accepted samples to correct for the discrepancy between observed and simulated summaries), but with far fewer user-chosen tuning knobs.

A key advantage of NPE in practice is that it “just plugs in” on top of whatever representation of the data is convenient. Given an expressive enough network, the same machinery can operate on hand-crafted summary statistics or on features extracted by a separate deep network, without needing to specify distance functions, tolerance thresholds, kernel shapes, or regression corrections as in ABC. This substantially reduces the amount of problem-specific hand-holding required from the user: once a simulator is in place and a prior is defined, NPE can be trained in a largely standardized way across very different population genetic applications. The trade-off is that, unlike some formulations of ABC, current NPE methods generally do not come with the same strong asymptotic guarantees of consistency for arbitrary choices of architecture and training scheme, and are instead justified empirically and by approximation arguments.

Importantly, we view the use of summary statistics as a central and enduring use case for NPE, rather than merely a stepping stone toward fully end-to-end models. When the summaries are interpretable and grounded in classical population genetic theory (e.g., site frequency spectra, LD statistics, etc.), NPE inherits those interpretability benefits while providing a fast, amortized posterior approximation. Because NPE learns a parametric approximation to the posterior, once trained it can evaluate posteriors for new datasets in milliseconds, making it particularly well-suited to settings where a small number of theoretically motivated summaries are available but inference must be repeated many times (e.g., across genomic windows, bootstrap replicates, or multiple populations).

In this study, we demonstrate that this approach—neural posterior estimation—offers incredible promise for the population genetics community by allowing the “best of all worlds” for simulation-based inference: interpretable and theoretically grounded summary statistics when desired, end-to-end feature extraction when needed, and in both cases rapid, amortized posterior evaluation with well-calibrated uncertainty. We illustrate how NPE can be used with both classical summaries and neural feature extractors (e.g., CNNs) that ingest genotype matrices directly, and how these components can be expressively composed so that feature extraction networks summarize the data before posteriors are estimated. In what follows, we demonstrate the utility of NPE in a number of population genetic settings, including recombination rate estimation and inference of demographic parameters: we show that learned posteriors are both rapidly generated, well-calibrated, and in some cases more accurate than those produced by currently available techniques.

## Methods

To date, the most popular approaches to simulation-based inference in population genetics are Approximate Bayesian Computation (ABC) and supervised machine learning approaches. A general schematic of these workflows is show in Figures 1a and 1b. Both of these approaches share a common strategy of defining a prior distribution *p*(*θ*) on the parameters of a model *θ*, and then using a simulator to generate data *x*^′^ conditional upon a draw from the prior; e.g. simulating from *p*(*x*^′^, *θ*) = *p*(*x*^′^|*θ*)*p*(*θ*). However, from that point in the procedure the two approaches diverge. ABC generates draws from the posterior p(*θ*|x) via approximate rejection sampling: by first reducing the simulated data *x*^′^ to a summary *y*^′^, and accepting the corresponding draw of *θ* if *ϕ*(*y, y*^′^) < *ϵ* (where *ϕ* is some measure of discordance between simulated and observed summary statistics, and *ϵ* is an arbitrary threshold). In practice the ABC rejection step may involve post-processing by regression or a more complex model to reduce bias and increase statistical efficiency. In contrast, the supervised machine learning approach aims to train a heavily parameterized model—e.g., an artificial neural network (ANN)—on the simulated, unreduced data to predict the value of *θ* which generated it. Through training of the ANN, the model can automatically create informative features from the raw data, in a process that is generally called automated feature extraction, rather than depend on the user’s choice of summaries to produce *y*^′^. The trained model applied to the observed data results in a point estimate of *θ* (e.g. Adrion et al., 2020a).

**Figure 1:**
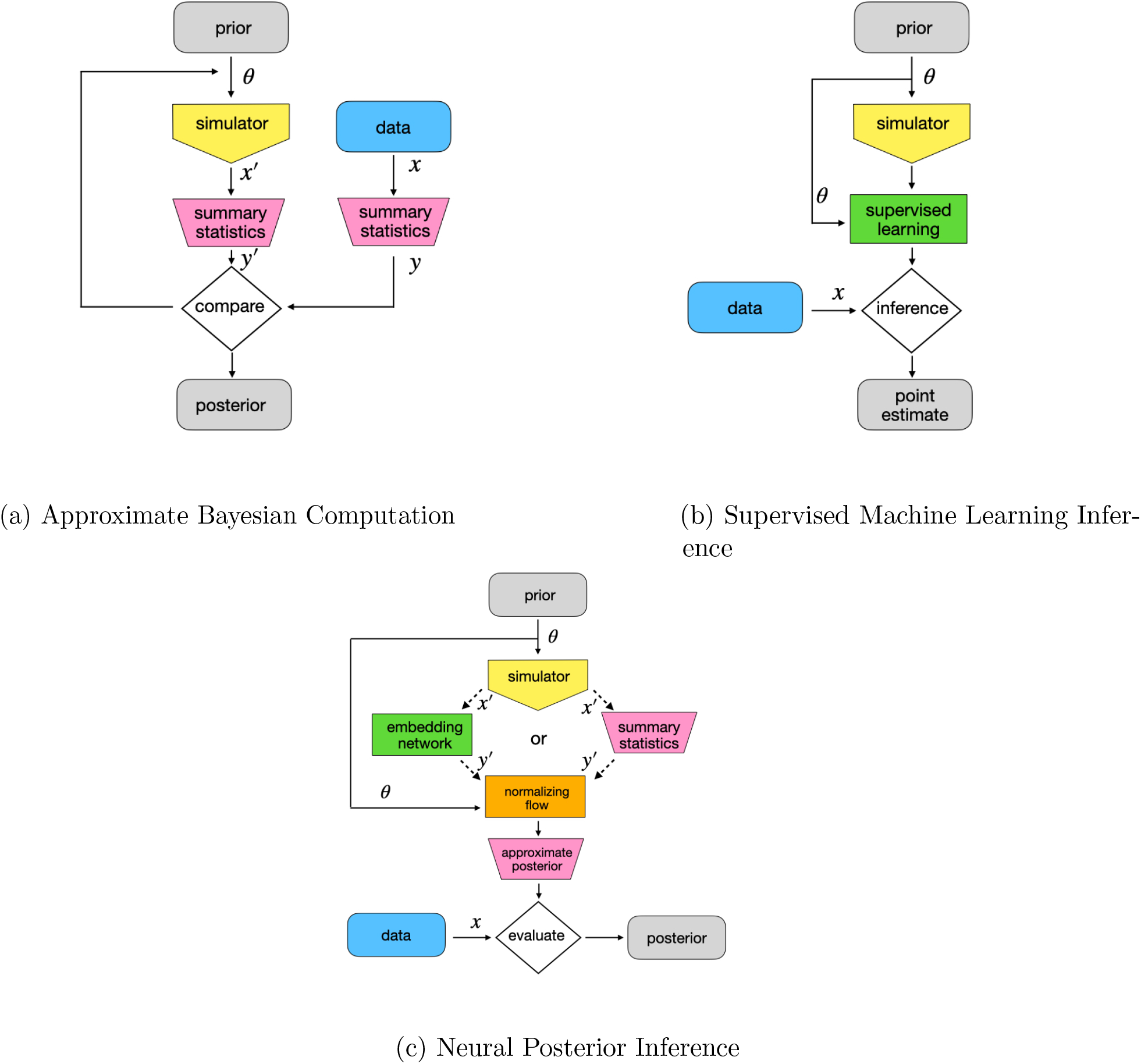
Schematic workflows comparing strategies of simulation-based inference in population genetics. Common workflows currently are Approximate Bayesian Computation (ABC; Fig. 1a) and Supervised Machine learning (Fig. 1b). Here we apply neural posterior inference (Fig. 1c) for inference in population genetics.

Here we apply a recently developed method for simulation-based inference of the posterior distribution of the parameters given the data, neural posterior estimation (NPE; Figure 1c that combines elements of ABC and machine learning). NPE also uses samples from *p*(*x*^′^, *θ*), but feeds these through a particular form of deep neural network designed for density estimation known as a *normalizing flow* that approximates this joint distribution. After sufficient training, the NPE model is conditioned on the observed data (or summaries thereof) to directly approximate the posterior *p*(*θ*|*x*). Like the supervised machine learning approach, NPE can operate on the original data rather than on summary statistics, potentially using an *embedding network* of arbitrary complexity prepended to the normalizing flow—however, the output is a distribution over plausible values of *θ* rather than a point estimate.

### Neural Posterior Estimation Overview

Neural posterior estimation (NPE) provides a flexible way to learn the posterior distribution *p*(*θ* | *x*) directly from simulated pairs (*θ, x*) using deep neural networks designed for density estimation. Whereas ABC relies on approximate rejection sampling and supervised machine learning produces point estimates, NPE learns an explicit, differentiable mapping that transforms a complex posterior distribution over *θ* into a simple latent probability distribution and vice versa. This is accomplished using *conditional normalizing flows*—invertible neural networks that implement learned changes of variables conditioned on the data (Rezende and Mohamed, 2015; Papamakarios et al., 2021).

Normalizing flows were introduced by Rezende and Mohamed (2015) and have since been extended to conditional density estimation for simulation-based inference (Papamakarios and Murray, 2016; Lueckmann et al., 2017; Greenberg et al., 2019); see Papamakarios et al. (2021) for a comprehensive review and Cranmer et al. (2020) for a broader overview of simulation-based inference methods. The central idea is to introduce a latent variable z with a tractable density (typically a standard multivariate Gaussian) and to learn an invertible, differentiable transformation

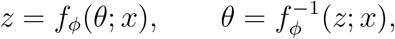

parameterized by network weights *ϕ*. Because *f*_*ϕ*_ is invertible, the density of *θ* conditional on *x* can be evaluated via the change-of-variables formula (Rezende and Mohamed, 2015; Papamakarios et al., 2021). Specifically, the approximate posterior density is

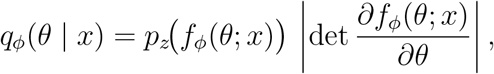

where *p*_*z*_ is the base density (e.g. a standard Gaussian) and the second factor is the absolute determinant of the Jacobian of the transformation. In practice, normalizing flows accomplish this by composing a sequence of simple, invertible transformations whose Jacobians are efficiently computable. In our implementation we use the `sbi` package (Tejero-Cantero et al., 2020) to construct a masked autoregressive flow whose elementwise transformations are monotone rational-quadratic splines (Durkan et al., 2019).

Training minimizes the expected negative log-posterior over the joint simulated distribution,

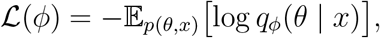

which is estimated by averaging over mini-batches of simulated (*θ, x*) pairs (Papamakarios and Murray, 2016; Lueckmann et al., 2017; Greenberg et al., 2019). Through this training on simulated (*θ, x*) pairs, the network learns how the posterior distribution *p*(*θ* | *x*) varies with the data, as minimizing this loss function is equivalent to minimizing the expected Kullback–Leibler divergence between the learned posterior *q*_*ϕ*_(*θ* | *x*) and the true posterior *p*(*θ* | *x*) averaged over datasets *x* generated by the simulator.

A key advantage of NPE in population genetic applications is that the input *x* is not restricted to hand-crafted summary statistics. Instead, *x* may consist of raw genetic data, image-like encodings of alignments, or summaries produced automatically by an *embedding network φ*. Such embedding networks—CNNs, RNNs, transformers, or other architectures appropriate to the data modality—learn informative low-dimensional representations from the raw data. These learned features are then provided to the normalizing flow, which maps them to an approximate posterior over *θ*. This compositional structure allows NPE to implement *end-to-end learning*: feature extraction and posterior estimation are learned jointly within the same model.

Training proceeds by optimizing the parameters of both the embedding network and the normalizing flow using stochastic gradient descent. All transformations are differentiable, enabling the use of automatic differentiation to compute gradients without resorting to MCMC, rejection sampling, or other sampling-based approximations. Once trained, the model provides amortized inference: at inference time, given new observed data *x*_obs_, posterior samples are generated by drawing *z* ∼ *p*_*z*_ and applying the inverse transformation 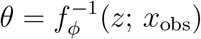, while the posterior density at any point *θ* can be evaluated directly via the change-of-variables formula above. No further simulation, optimization, or MCMC sampling is required—a single forward pass produces the full approximate posterior distribution. Figure 2 illustrates this inference workflow.

**Figure 2:**
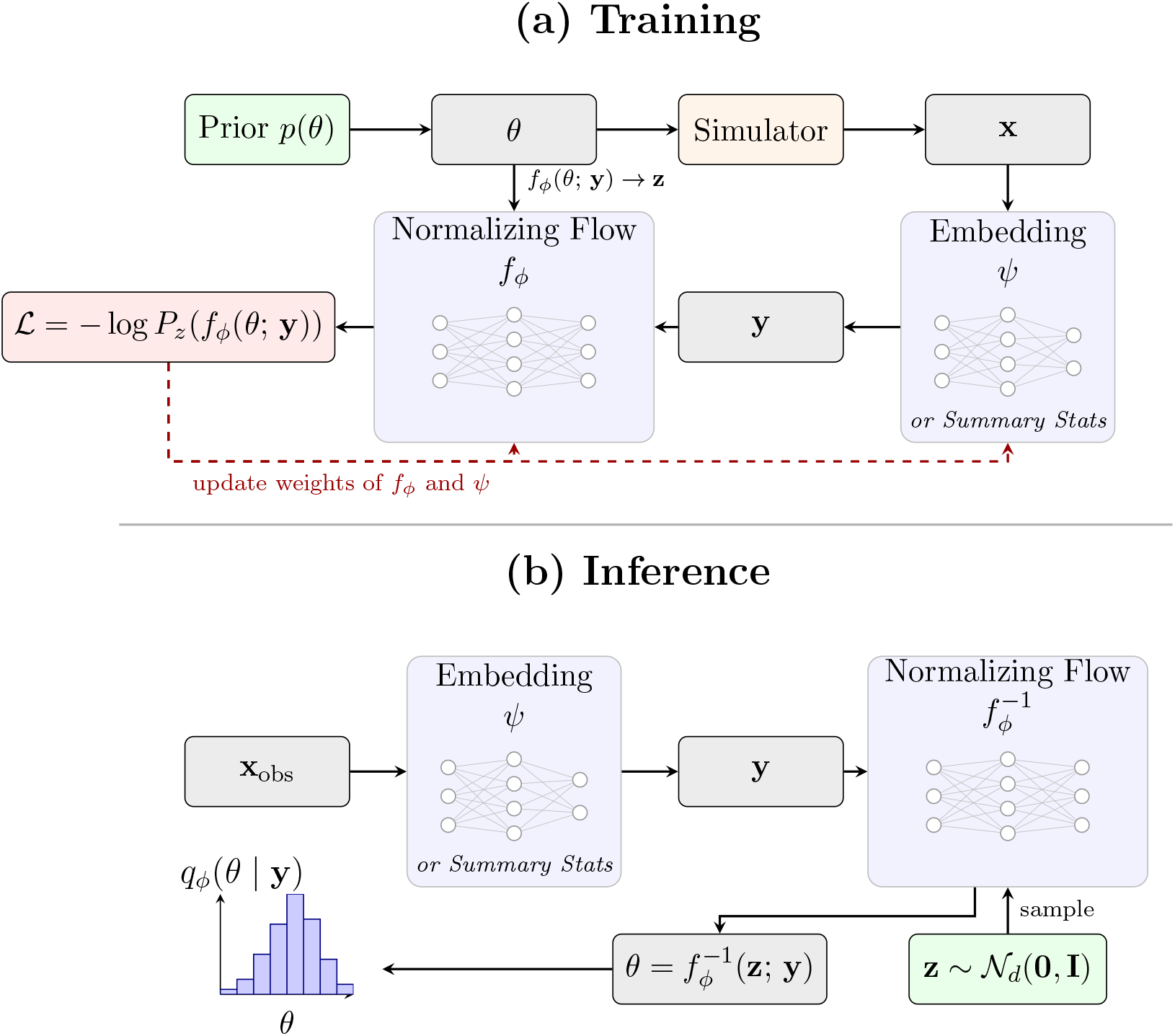
Schematic of neural posterior estimation showing the **(a)** training and **(b)** inference stages. **(a)** During training, parameters *θ* are drawn from a prior *p*(*θ*) and used to generate simulated data **x** via a population-genetic simulator. The data are then *either* passed through a learned embedding network *φ or* reduced to user-chosen summary statistics to produce a summary vector **y**. The normalizing flow *f*_*ϕ*_ maps *θ* to a latent variable **z**, conditioned on **y**; the loss *ℒ* = − log *P*_*z*_(*f*_*ϕ*_(*θ*; **y**)) evaluates the transformed parameters under the base density and is minimized by updating the weights of both *f*_*ϕ*_ and *φ* (dashed arrows). **(b)** At inference, observed data **x**_obs_ are embedded to produce **y**, samples **z** ∼ 𝒩_*d*_(**0, I**) are drawn from the base distribution, and the inverse flow yields posterior samples 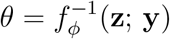, giving the approximate posterior *q*_*ϕ*_(*θ*|**y**). Note that **z** has the same dimension as *θ* (the parameter vector), not **y**.

### NPE Workflow

We have developed a `snakemake` pipeline targeted towards posterior estimation from genotype data, using arbitrary embedding networks. Two constraints on simulation-based inference using genome-wide population genetic data are that (a) genome-scale simulations are very costly to produce, and (b) the “raw” data (genotype arrays) are too large to fit in GPU memory. Hence, we take an amortized approach, where an NPE model (embedding network attached to normalizing flow) is trained on shorter simulated sequences then applied over the observed data in windows, to generate a sequence of posteriors. Local posteriors may be sufficient for the task at hand (e.g. recombination rate prediction, sweep detection) or may be averaged (or otherwise combined) to produce global estimates (e.g. for demographic inference). The workflow prepares training and testing datasets by simulating tree sequences with `msprime` (Baumdicker et al., 2022), performs preprocessing (as appropriate; see below), trains NPE estimators, estimates model parameters across genomic intervals, and subsequently summarizes results from estimation. Our workflow and all associated code can be found at https://github.com/kr-colab/popgen-npe. We use normalizing flow architectures from the `sbi` Python package Tejero-Cantero et al. (2020), and use a modular design such that arbitrary neural networks may be used to generate embeddings. We note that for some applications, a fully amortized approach may be statistically inefficient (in the sense that training examples simulated from the prior are too sparsely scattered across parameter space); in which case sequential refinement of the posterior estimator via importance sampling may be necessary (Tejero-Cantero et al., 2020).

### Population Genetic Tasks

Throughout this manuscript we apply NPE to a variety of common population genetic tasks, highlighting the benefits of NPE and contrasting its performance to established methods both from simulation-based methods and from probabilistic methods when appropriate. Many of the methods are shared among tasks and outlined in separate sections below.

#### Coalescent Simulation

All simulation for training and testing was performed using `msprime` (Baumdicker et al., 2022). Specific parameterizations were task dependent and details are given in the subsequent sections. For estimation of demographic parameters, demographic models are either written in `demes` (Gower et al., 2022) from scratch or adapted from a model from the `stdpopsim` catalog (Adrion et al., 2020b; Lauterbur et al., 2023), by letting a subset of the parameters (i.e. those targeted in estimation) be replaced by a prior sample.

#### Embedding Networks and Summary Statistics

Throughout the manuscript we tried a number of different embedding network architectures, referred to as *φ* in Figure 2, for automated feature extraction from raw genotype data and in places contrast performance with hand-designed collections of summary statistics.

For recombination rate estimation in Section *Recombination rate estimation*, we set *φ* to the recurrent GRU architecture of Adrion et al. (2020a) reimplemented in `PyTorch` to be compatible with our NPE workflow. This network takes as input the genotype matrix as well as a vector of positions encoding the locations of SNPs. The genotype matrix is passed through a bi-directional GRU layer, followed by a series of dense and dropout layers. The position input vector is separately passed through a dense layer, and then concatenated with the dropout layer output from the genotype branch of the network.

For inference of demographic parameters in Section *Inference of Population Bottleneck Parameters*, we used an exchangeable convolutional neural network (CNN) architecture modeled after Chan et al. (2018) and similar to the discriminator network implemented in `pg-gan` (Wang et al., 2021). Genotype and relative SNPs position information for input to these networks were prepared using the feature extractors in `dinf` (Gower et al. (2023)).

For inference of effective population size histories in Section *Inference of Population Bottleneck Parameters*, we again contrasted different choices of embedding neural networks with a summary statistic approach. Three DNN architectures were compared: 1) an RNN, 2) the exchangeable CNN architecture from Chan et al. (2018), and 3) a bespoke DNN architecture called `SPIDNA` from Sanchez et al. (2020). `SPIDNA` is designed specifically for genome sequence data, and incorporates haplotype permutation invariance while capturing long range dependencies between SNPs and variation in SNP density among regions. Due to its sophisticated design, training NPE with embedding `SPIDNA` demands the most computing power. In addition we examine performance in NPE inference when using either the SFS or LD statistics alone, for larger sequence lengths where end-to-end learning (e.g. from the raw genotypes) is infeasible.

For inference of effective population size histories in Section *Priors for Historical Population Size Inference*, we used a combination of linkage disequilibrium (LD) and site frequency spectrum (SFS) summary statistics as input features for neural posterior estimation. Simulated genomic regions of 2 Mb were divided into non-overlapping 1 Mb windows, and summary statistics were computed for each window and then averaged across windows. For LD, we computed the squared correlation coefficient (*r*^2^) between pairs of SNPs using the Rogers-Huff estimator (Rogers and Huff, 2009). SNP pairs were subsampled at gap sizes following powers of two, and *r*^2^ values were binned into 19 logarithmically-spaced distance classes ranging from 10^2^ bp to the window length. The mean *r*^2^ was computed for each distance bin. For the SFS, we computed the polarized (unfolded) site frequency spectrum for each window using `tskit` (Jeffery et al., 2026), excluding monomorphic site classes. With 25 diploid individuals (50 haplotypes), this yielded 49 SFS bins after excluding the zero and fixed classes. The final feature vector concatenated the 19 mean *r*^2^ values with the 49 mean SFS values, resulting in 68 summary statistics per simulation that were passed directly to the neural density estimator via a summary statistics embedding layer.

#### Comparisons to moments

In Section *Inference of Population Bottleneck Parameters* we directly compare point and uncertainty estimates derived from NPE to those from the popular composite likelihood approach `moments` (Jouganous et al., 2017). As `moments` takes the SFS as input, we compare its performance to NPE by using the normalized SFS alone as the summary statistics for input to the normalizing flow (see above). Optimization of `moments` was performed by an exhaustive grid search within the bounds of the prior. Uncertainty estimation using `moments` was performed using the Godambe information matrix routine with `eps` = 0.05.

#### Comparisons to Approximate Bayesian Computation

In Section *Inference of Population Bottleneck Parameters* we additionally compare NPE estimators to estimates based on approximate Bayesian computation (ABC). In this comparison we utilized the same set of training simulations used to train the NPE, but performed ABC rejection sampling (*c*.*f*., Pritchard et al., 1999), where the top 1% quantile of closest simulations was retained as the posterior sample. We used three summary statistics for the ABC as our summary vector: *π*, the number of segregating sites, and Tajima’s D. The joint posterior distribution is visualized using a kernel density estimate of the accepted simulation parameter values.

#### Comparisons to MSMC2

In Section *Inference of Historical Population Sizes in a One-Population-Model* we compare NPE-derived posteriors to estimates from MSMC2 (Schiffels and Durbin, 2014), a widely used pairwise sequentially Markovian coalescent method for inferring historical effective population sizes. For each demographic scenario, we subsampled 10 diploid individuals (20 haplotypes) from the simulated tree sequences used to evaluate the neural network approaches, converted the phased genotype data to MSMC2’s `multihetsep` input format (filtering multiallelic sites), and ran MSMC2 with a fixed recombination-to-mutation ratio of 1.0 (corresponding to equal per-base rates of *r* = *µ* = 1.5 × 10^−8^), 18 EM iterations, 20 threads, and the time segment pattern `1*2+25*1+1*2+1*3`. When MSMC2 did not converge to produce a final output, we selected the iteration with the highest log-likelihood from the intermediate results. Coalescence rates were converted to effective population sizes as *N*_*e*_ = 1/(2*λµ*), and scaled times were converted to generations by dividing by the mutation rate. MSMC2 estimates were overlaid as dashed lines on the neural posterior credible interval plots for each scenario.

#### Application to *Drosophila melanogaster*

We analyzed whole-genome sequence data from 19 haploid genomes (derived from inbred lines), including 9 sampled from France (FR) and 10 from Cameroon (CO), which were taken from the published DPGP2 collection (Pool et al., 2012). Prior to analysis, we filtered out genomic regions in which any sample contained missing data, ensuring that all downstream inference was based on complete genotype calls. Demographic inference was conducted under a modified Li and Stephan out-of-Africa model (Li and Stephan, 2006), which incorporates isolation with migration and exponential growth in the European lineage. This model includes seven free parameters: the ancestral population size, the effective sizes of the African and European populations following the split, the contemporary European effective size, the time of population divergence, and asymmetric migration rates between the European and African populations. For all parameters, we use uniform priors on a log scale (10^3.5^ to 10^6.5^ for population sizes; 10^3^ to 10^6^ for the split time; and 10^−8^ to 10^−3^ for migration rates).

We employed neural posterior estimation (NPE) to infer posterior distributions of these demographic parameters, using a recurrent neural network embedding trained on 300 Kb non-overlapping windows across chromosome arm 2L (each window containing approximately 6,000 SNPs). In each simulation, the population-scaled mutation rate *θ* was fitted assuming a per-base mutation rate of 0.45 × 10^−8^, which accounts for the observed rate of missing data and corresponds to a correction from the baseline rate of 0.54 × 10^−8^ reported by Schrider *et al*. (2013). This rescaling ensures that simulated nucleotide divergence between African and European samples matches the observed value, and is analogous to the standard practice of fitting *θ* in site-frequency spectrum–based inference. The recombination rate was treated as a nuisance parameter and integrated out under a uniform prior ranging from 0.5 × 10^−8^ to 3 × 10^−8^ per base per generation. Input genotype features were unpolarized, with the minor allele at each site taken as the “derived” state.

To evaluate model fit, we performed posterior predictive checks by generating new sequence data for each window from the posterior distribution using `msprime`, and then calculated summary statistics (*π, F*_*ST*_, Tajima’s *D*). Posterior predictive distributions closely matched the observed statistics, indicating that the model adequately captured the data (Fig. 11). Moreover, systematic genomic variation in demographic parameter estimates was reflected in the posterior distributions across windows (Fig. S12). Finally, we averaged posteriors across windows to obtain global demographic parameter estimates, which are summarized in Fig. 10.

## Results

### Recombination Rate Estimation

Before tackling high-dimensional problems, we start with a 1-dimensional problem—inferring a uniform recombination rate. Our particular goal here is to show that NPE derived posteriors are well-calibrated and computationally efficient.

Previously, Adrion et al. (2020a) introduced a recurrent neural network (`ReLERNN`) that learns recombination rates from genotype data. The `ReLERNN` architecture consists of a bidirectional recurrent (GRU) layer that takes raw genotypes as input, followed by a series of fully-connected layers which end in a scalar output (the predicted recombination rate). Adrion et al. (2020a) showed that parametric bootstrapping provides well-calibrated uncertainty estimates: given a predicted recombination rate, genotype data is simulated via `msprime` (Baumdicker et al., 2022) and the trained `ReLERNN` model is applied to each synthetic dataset, generating a distribution of bootstrap replicates from which confidence intervals may be readily calculated. However, when the trained network is used across a large number of genomic windows (as would be the typical application, to build a recombination rate map) this procedure becomes expensive as a new set of simulations must be generated for every window. An alternative approach based on NPE is appealing as the cost of genotype simulation is paid up front.

For the current comparison we reimplemented the `ReLERNN` architecture within our pipeline. For training, we used coalescent simulations of ten diploids over a sequence of length 1Mb with an effective population size of 10^4^ and a variable per-base mutation rate uniformly drawn from [0.66 × 10^−8^, 1.33 × 10^−8^]. For each simulation, the recombination rate *r* was sampled from a uniform [0, 10^−8^]. We trained `ReLERNN` with 20,000 simulations with mean squared error loss against true r, and applied the parametric bootstrap procedure described above to get confidence intervals for a test set of 2,000 examples (using 1,000 bootstrap replicates per example).

To generate a comparable set of posteriors with NPE, we first froze the trained `ReLERNN` network without dropout and removed the last fully-connected layer (that collapses a latent space of size 64 to a scalar output). With this “decapitated” pre-trained network, we generated 64-dimensional summary statistics for each input sequence. We used these learned **embeddings** of the genetic data to train an NPE model, then generated posterior distributions for the same test set that was used for parametric bootstrapping. Finally, we compared the NPE and bootstrapped uncertainty estimates by measuring confidence/credibility interval coverage (Figure 3) and concentration (Figure 4) across a range of interval widths. We found that both methods produced similarly well-calibrated intervals; however, the NPE procedure was far less computationally expensive, as it does not require generating novel simulations per prediction. These results show that NPE can be used to learn posteriors for recombination rates with comparable calibration but large computational savings in comparison to parametric bootstrapping.

**Figure 3:**
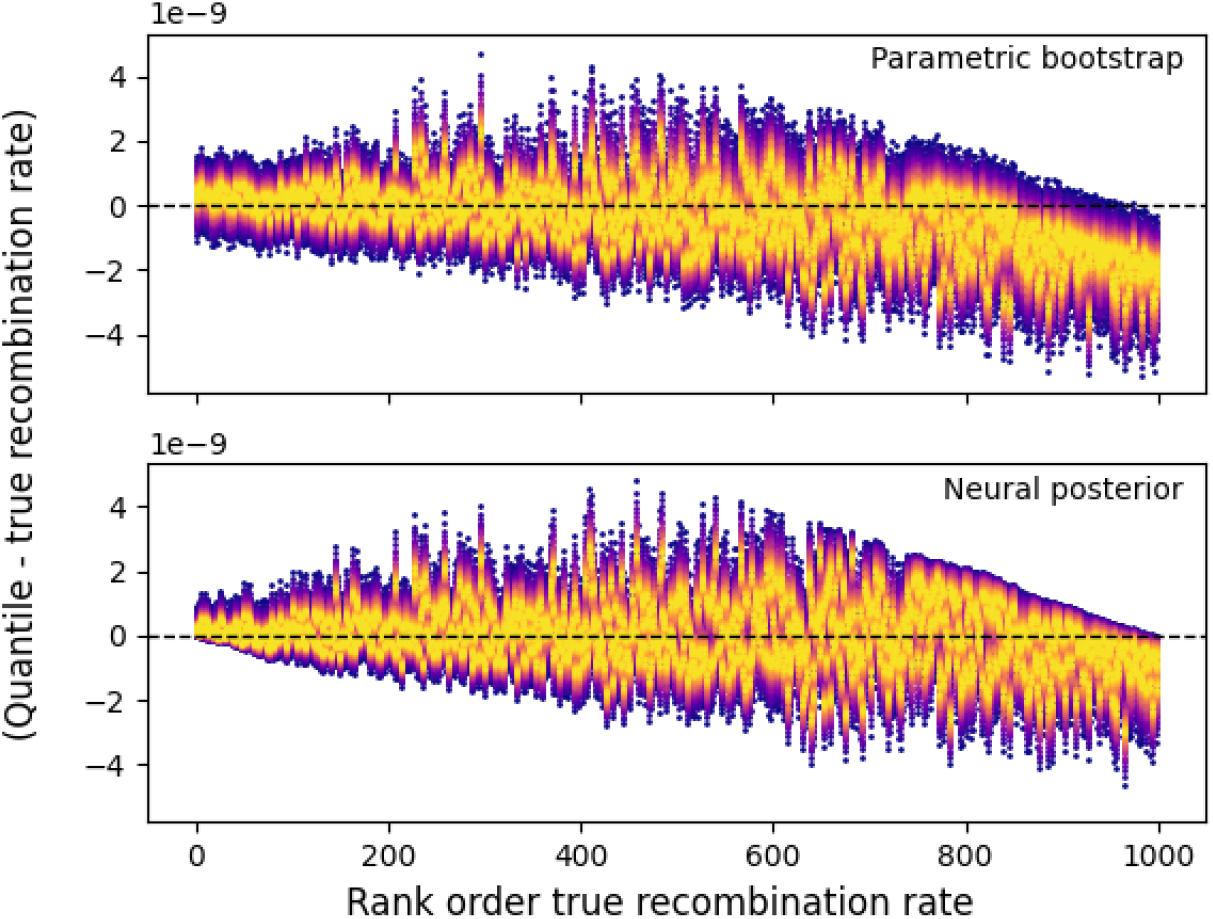
Posterior intervals for recombination rate estimation. Each column represents one of 1,000 test simulations, ranked by true recombination rate (x-axis). Colored bands show the credible/confidence interval quantiles (inner to outer: 10% to 95%, in increments of 10%) minus the true recombination rate, for parametric bootstrapping (top) and neural posterior estimation (bottom). Well-calibrated intervals should be centered on the dashed zero line, with widths that reflect the true uncertainty. Both methods produce similarly centered intervals; the NPE intervals are slightly narrower, indicating comparable calibration with greater concentration.

**Figure 4:**
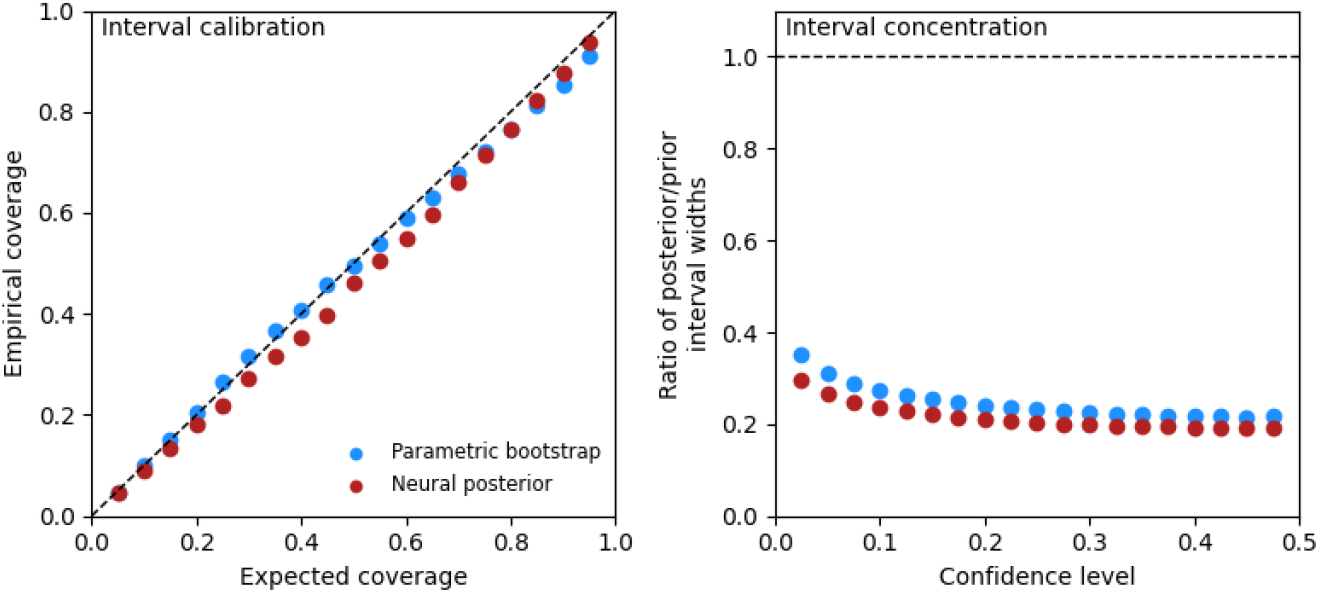
Posterior coverage and concentration for recombination rate estimation. Left: empirical coverage versus expected coverage for parametric bootstrap confidence intervals (blue) and NPE credible intervals (red). The dashed diagonal indicates perfect calibration; both methods closely track this line, indicating well-calibrated uncertainty estimates. Right: interval concentration, measured as the ratio of posterior (or bootstrap) interval width to the prior width, plotted against confidence level. Values well below 1 (dashed line) indicate that the posteriors are substantially more concentrated than the prior. Both methods achieve similar concentration, with NPE intervals slightly narrower at most confidence levels.

To investigate the impact of prior choice on the efficiency of training, we performed an additional analysis where we trained NPE networks on three different priors and increasing numbers of simulated instances, then generated posterior predictions over the same test set, simulated from a uniform distribution on [0, 3 × 10^−8^]. When given a non-informative but overly wide prior, the NPE required more instances to train, but ultimately converged to an equivalent posterior as the NPE trained on the true prior (Figure S1, middle). In contrast, when given a prior with the same support as the true prior but a “overly-concentrated” shape (i.e. with a mode at 1.5 × 10^−8^), the NPE trained much more efficiently but produced too-narrow intervals across the test set, as expected (Figure S1, right). We next move on to a simple two-dimensional problem: inferring the timing and magnitude of a bottleneck event.

### Inference of Population Bottleneck Parameters

Our next task is to infer the timing and the intensity of a bottleneck event. For this task we use *Arabidopsis thaliana* as an example, building upon the South Middle Atlas African demographic model from the `stdpopsim` catalog (Huber et al. (2018); Adrion et al. (2020b); Lauterbur et al. (2023)).

Training simulations consisted of 100 Kb of simulated data from chromosome 1 using the recombination map of Salomé et al. (2012). For our training set for NPE, we simulated 20,000 tree sequences with *T* (bottleneck time in generations from present divided by twice the ancestral population size) and *ν* (nu; the ratio between the current population and the ancestral population sizes) sampled from uniform distributions Unif(0, 1.5) and Unif(0, 1) respectively.

As a baseline comparison we used the `moments` package (Jouganous et al. (2017)) to infer the bottleneck parameters from the same simulated data. `moments` uses a composite likelihood approach to infer the bottleneck parameters from the site-frequency spectrum (SFS) of a sample and utilizes the Godambe information matrix to estimate the uncertainty of the parameter estimates (which is asymptotically correct, despite the model mis-specification intrinsic in composite likelihood). Throughout this section, we compare the performance of `moments` to NPE-derived estimates and posteriors using three different embedding approaches: 1) using the SFS as input as a form of summary statistics (i.e. without using the embedding network), 2) using a 1D convolutional neural network (CNN) to embed raw genotypes generated through simulation, and 3) using a recurrent neural network (RNN). We note that using the SFS as input to NPE amounts to a direct comparison to `moments` as the same information is available to both methods. As an additional point of comparison for this task we did ABC rejection sampling of our NPE training simulations, using a simple set of summary statistics known to be informative for population bottlenecks (*π*, the number of segregating sites, and Tajima’s D). We retained the closest 1% of simulations as our posterior sample.

An important starting point is to compare the `moments` derived likelihood surface to the NPE-derived posteriors. In Figure 5 we show comparisons of the NPE posteriors to the `moments` likelihood surfaces and the ABC-derived posterior distribution for three independent points in parameter space (given on rows). Time of the bottleneck (*T*) is shown on the y-axis and the intensity of the bottleneck (*ν*) is shown on the x-axis. True parameter values are indicated by a red dot. For the NPE posteriors, we show the results from three different embedding approaches labeled npe-cnn, npe-rnn, and npe-sfs, respectively. As can be seen from the figure, the likelihood and posterior densities of *T* and *ν* show that the parameters are correlated in a non-linear fashion. This feature is problematic from the standpoint of likelihood-based quantification of uncertainty, as canonical confidence intervals based on Fisher information, including Godambe information, are not appropriate as they assume a Gaussian form. Qualitatively, all methods contain the true parameter values within their respective areas of high density. However, there is some variation among the NPE embedding approaches.

**Figure 5:**
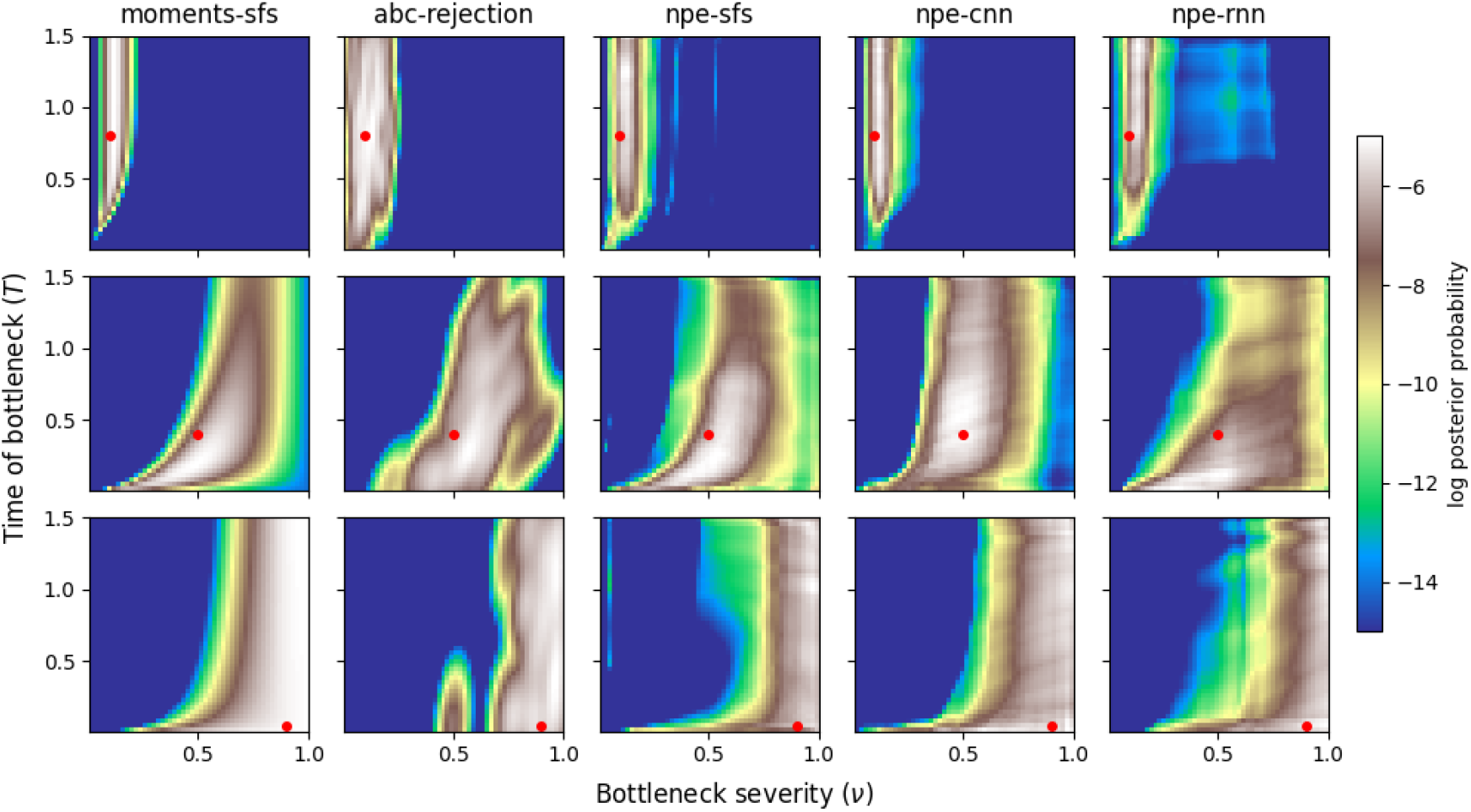
Comparison of NPE posteriors to ABC and moments likelihood surfaces. Here we show comparisons of the NPE posteriors to the ABC-derived posteriors and the `moments` likelihood surfaces for three independent points in parameter space (given on rows). Time of the bottleneck (*T*) is shown on the y-axis and the intensity of the bottleneck (*ν*) is shown on the x-axis. True parameter values are indicated by a red dot. For the NPE posteriors we show the results from three different embedding approaches labelled npe-cnn, npe-rnn, and npe-sfs. The likelihood surface for `moments` is scaled to match the curvature of the Godambe Information Matrix at the mode and normalized by the total mass within the prior bounds, to make it comparable to the posterior densities. Colors indicate the likelihood/posterior density.

We can further characterize the performance of the NPE approach by looking at the point estimates and their associated uncertainty. In Figure 6 we show scatterplots of the NPE-derived point estimates compared to the `moments` and ABC point estimates for each of the three embedding approaches. As the NPE approach yields a distribution of point estimates, we show the mean of the posterior distribution as a point estimate, corresponding to the Bayes estimate. For each estimator we also show the estimated mean squared error (MSE) loss across the parameter space for each parameter (labeled MSE). As can be seen from the figure, the NPE-derived point estimates each yield a lower loss than the `moments` point estimates and equivalent to that from ABC, indicating that the NPE approach is able to accurately estimate the true parameter values. As an additional view of this comparison, Figure S2 shows boxplots of MSE loss for the NPE-derived point estimates compared to the `moments` and ABC point estimates across parameter space for *ν* and *T* (left and right panels, respectively). Again, the general trend here is that the NPE-derived point estimates yield a slightly lower loss than the `moments` point estimates.

**Figure 6:**
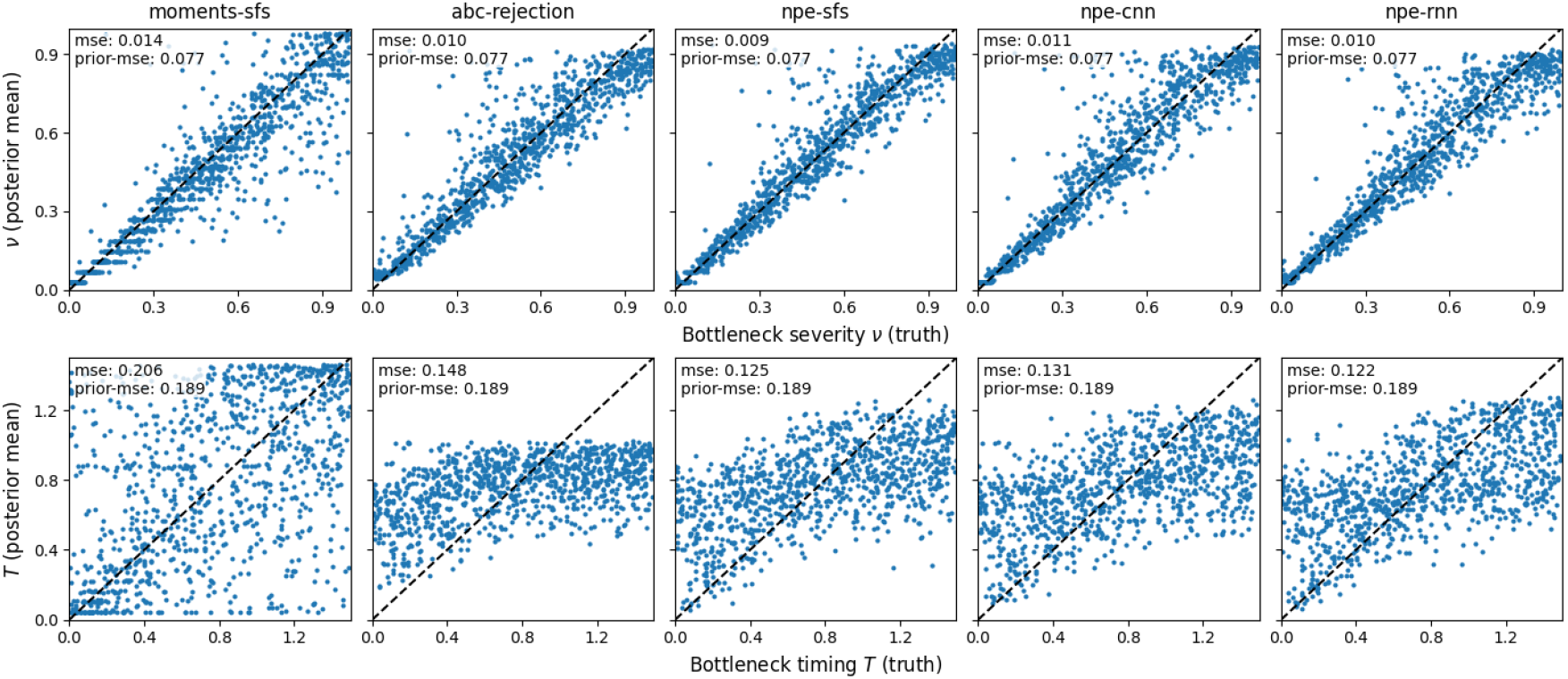
Comparison of NPE point estimates to moments and ABC point estimates. Here we show comparisons of NPE-derived point estimates to the `moments` and ABC point estimates across parameter space for *T* and *ν* (given on rows). Columns show the different embedding approaches: npe-cnn, npe-rnn, and npe-sfs. As the NPE approach yields a distribution of point estimates, we show the mean of the posterior distribution as a point estimate, corresponding to the Bayes estimate. For `moments`, we take the maximum likelihood estimate from a brute-force grid search within the prior bounds. Each facet additionally shows the estimated loss across the parameter space for each estimator (labeled MSE) as well as the loss relative to the prior mean (labeled prior-mse).

Ultimately our goal is to quantify uncertainty in the point estimates, rather than point estimation itself. To this end, we can compare the coverage of the estimated posterior across our embedding approaches to the coverage of Godambe information matrix-derived confidence intervals for `moments` as well as that from the ABC-derived posterior. In Figure 7 we show the coverage of the NPE-derived posteriors across our embedding approaches to the coverage of Godambe information matrix (GIM)-derived confidence intervals for `moments`. There are two major takeaways from this figure. First, the NPE-derived posteriors display excellent coverage for both *ν* and *T* . As with our recombination rate results, this is a strong indication that the NPE-derived posteriors are well-calibrated. Second, the `moments`-derived confidence intervals are less well-calibrated for *T*, in particular as *T* gets larger. As pointed to above in looking at the posterior surfaces, this is a results of the non-linear relationship between *T* and *ν*; GIM-derived confidence intervals are not able to capture the full complexity of the likelihood surface, and as a result the confidence intervals are too narrow.

**Figure 7:**
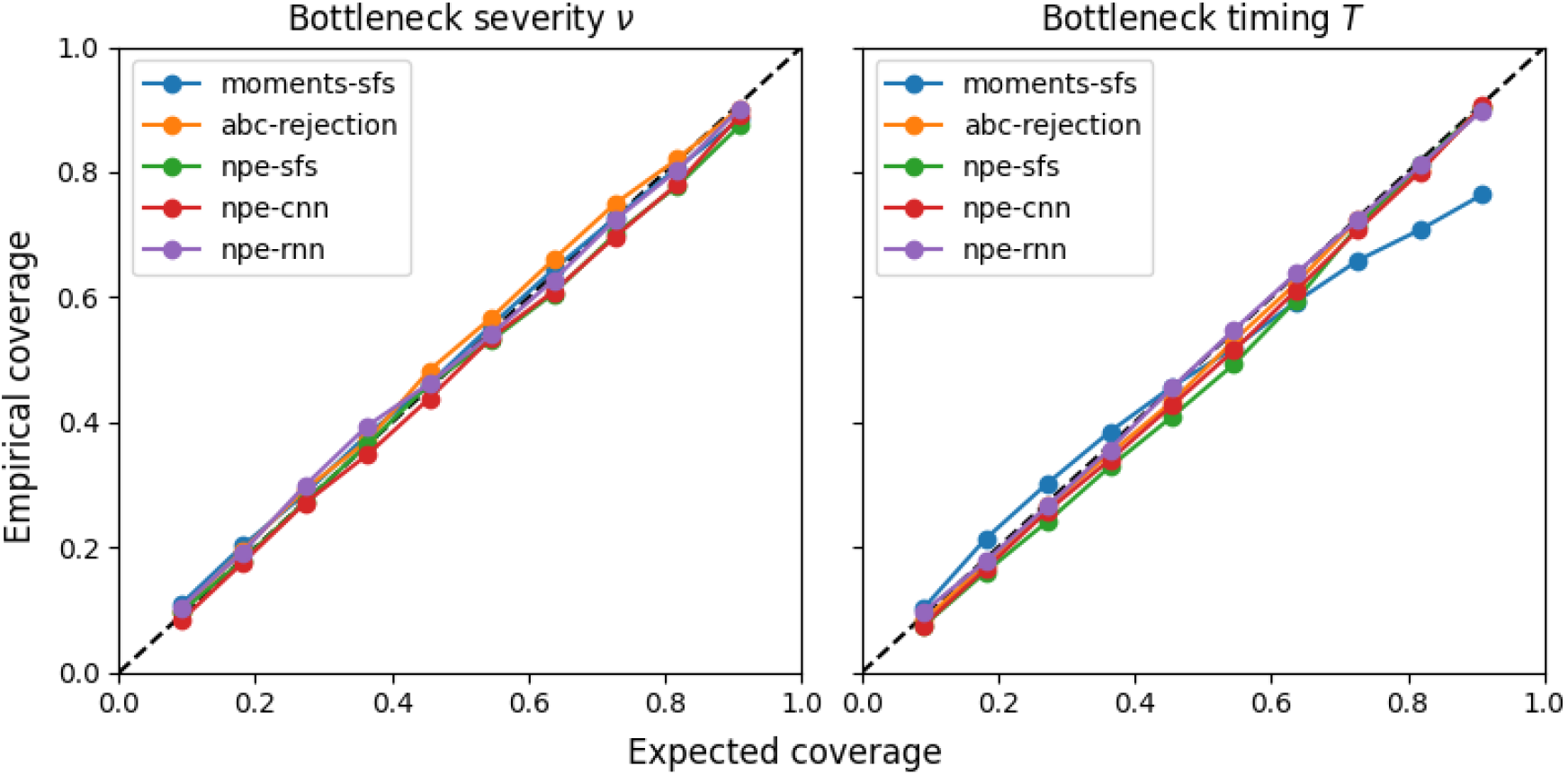
Comparison of NPE and ABC posterior coverage to `moments` confidence intervals. Here we show comparisons of the empirical and expected coverage of the NPE-derived and ABC-derived posteriors across our embedding approaches to the coverage of Godambe information matrix-derived confidence intervals for `moments`. The left and right panels show the coverage for *ν* and *T*, respectively. Labels are as in Figure 5.

As the NPE and ABC procedure depend on a prior distribution, and the `moments` approach is based on bounded optimization, an additional comparison of posterior concentration is warranted. Ideally, any Bayesian approach should yield a posterior distribution that is more concentrated around the true parameter values than the prior distribution. In Figure S3 we show the ratio of the interval widths of the prior and posterior distributions across for *ν* and *T* for each of the three embedding approaches as well as to ABC a comparison point. We compare this to `moments` by using as a prior the bounds on the optimization and considering as the posterior the interval width of the Godambe information matrix-derived confidence intervals. As can be seen from the figure, the NPE-derived posteriors are heavily concentrated and show a large reduction in the interval width compared to the prior for both parameters. Indeed for our choice of summary statistics, the NPE posterior is even more heavily concentrated than is the ABC-derived coverage. Again this is an indication that the NPE procedure is effectively learning the posterior distribution.

A caveat is that our simulated data is “small”, in the sense that we are only using 100 Kb of sequence per simulated instance, and both the `moments` likelihood surface and the learned NPE posteriors should improve with additional data. Indeed, repeating the experiment described above with simulated 1 Mb sequences demonstrates that this is the case (Figures S4 through S7), while the differences between estimators and embedding networks remain qualitatively similar. Notably, structural non-identifiability is maintained in the NPE posteriors regardless of the increased sequence length. For example, in the top and bottom rows of both Figures 5 (100 Kb) and S4 (1 Mb) the bottleneck timing is poorly identified, because the bottleneck severity is either so extreme or so weak that variation in the timing has little impact on the observed data. This intrinsic source of uncertainty is reflected in both sets of posteriors; whereas for “well-identified” intermediate scenarios (e.g. the middle row of Figures 5 and S4), the posterior concentrates for both parameters with increasing information.

### Inference of Historical Population Sizes in a One-Population-Model

Another task that we were keen on exploring is the inference of effective population size over time. This is a common task in population genetics, and it is often used to infer the population size history of a species modulo caveats about the relationship between the effective population size and the census population size. While specialized methods for this task exist (e.g., Li and Durbin, 2011; Schiffels and Durbin, 2014; Terhorst et al., 2017), we were interested in seeing if a neural posterior estimator could be used to infer the effective population size over time. Indeed, others have explored the use of Bayesian methods for inference of effective population size over time both with and without neural networks (Boitard et al., 2016; Sanchez et al., 2020).

To train the neural posterior estimator, we simulated 40,000 tree sequences with `msprime`. For each of the 40,000 tree sequences, we simulated histories with 15 population size changes for samples of size *N* = 20 individuals. Population size changes occurred on a fixed grid of 16 log-spaced time points from 10^2^ to 10^5^ with # log_10_ ≈ 0.2. Population sizes for each of the 15 time epochs were sampled independently from a log-uniform distribution with log_10_(*N*_*e*_) ∼ Uniform(2.0, 5.0), corresponding to effective population sizes ranging from 100 to 100,000 individuals. Training of the neural posterior estimator then followed as in the previous sections. We compared three different embedding networks: a CNN embedding network (as above), a RNN embedding network, and finally the SPIDNA embedding network from (Sanchez et al., 2020). We chose to include the SPIDNA embedding network as it was specifically designed for embedding population genetics data for the task of population size inference.

We benchmarked the performance of the neural posterior estimator for inferring effective population size over time under three demographic scenarios: 1) an abrupt change in population size, 2) a scenario of a power-law population growth with exponent 1/2, and 3) a scenario of power-law population decline with exponent 1/2. In each case, the trained neural posterior estimator was used to infer the effective population size over time. Figure 8 shows the results of this benchmark for each of our three embedding networks. Generally, the neural posterior estimator is able to recover the true population size history relatively well, with each of the three embedding networks containing the true population size within the 50% credible interval for the vast majority of the time epochs. Generally performance declines moving back in time, as the amount of information available decreases. This can be seen more clearly in Figure S8 where we have added a line showing the percentage of remaining lineages at each time point within the test simulation.

**Figure 8:**
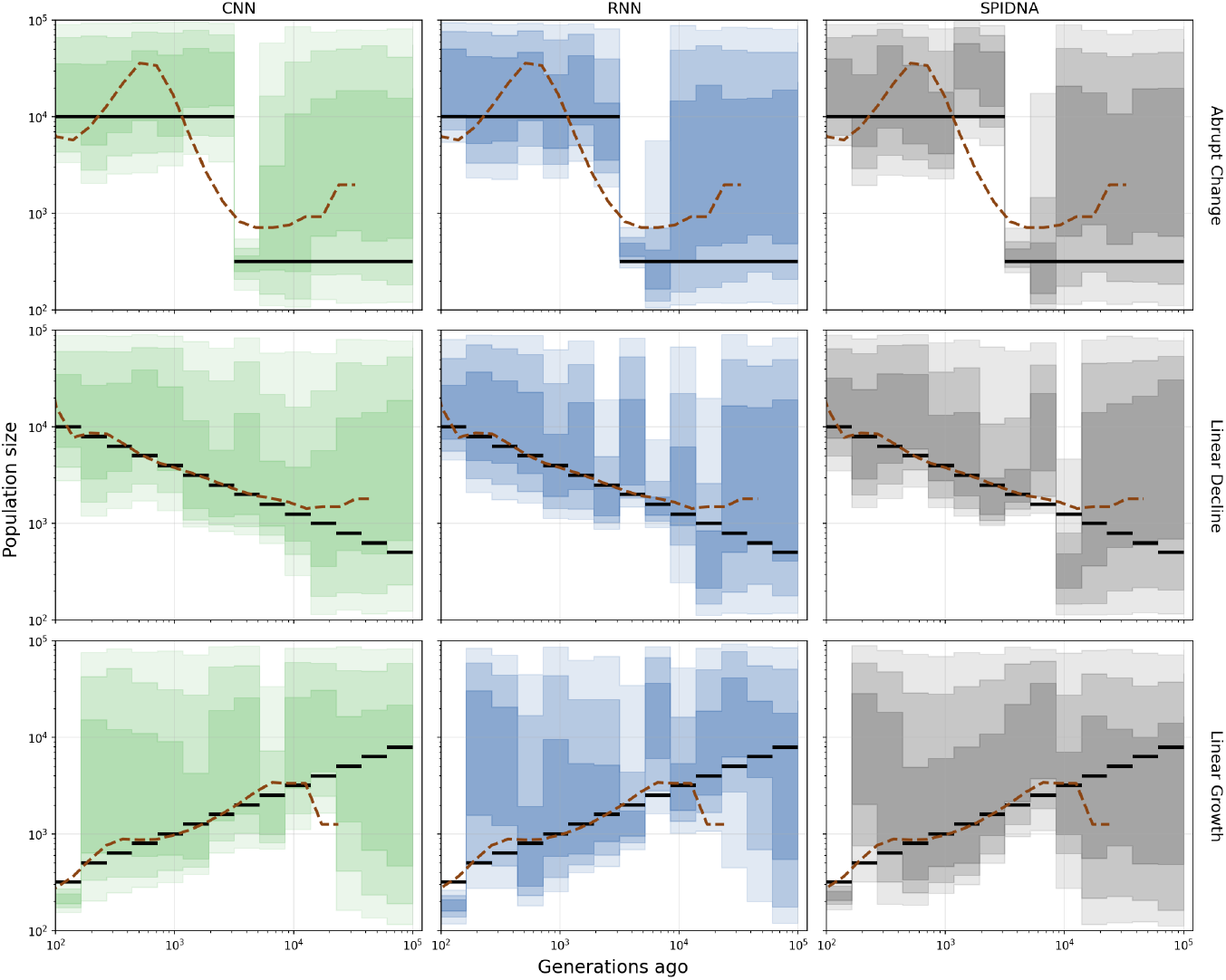
Benchmark of the neural posterior estimator on the task of inferring the effective population size over time using three different embedding networks. The three scenarios shown on rows are: 1) an abrupt change in population size, 2) population growth, and 3) population decline. The three embedding networks shown on columns are: a CNN embedding network, a RNN embedding network, and the SPIDNA embedding network from (Sanchez et al., 2020). For each plot we show the true population size history (black line), the inferred credible intervals (shaded regions) for the 15 time epochs, and the MSMC2 estimate (dashed line) as a classical baseline.

In Figure S9 we show the normalized credible interval (CI) width for each of the parameters estimated by the NPE for each of the scenarios and each of the embedding networks. Note that we estimated 15 population size parameters as well as the recombination rate. Normalized credible interval width was calculated as the 95% credible interval width divided by the geometric mean for population size parameters and by the posterior mean for the recombination rate parameter. The geometric mean was used for population sizes as the prior is log-uniformly distributed, where as the posterior mean for the recombination rate as its prior was drawn from a linear space. We see that there is some variation in the inferred CI width among embedding networks that differs across the considered demographic scenarios. There is a general tendency for CIs to grow wider for population size parameters that describe more ancient times, which we again interpret as the models reflecting less information as time moves towards the past.

Finally, in Figure S10 we show mean relative errors for aggregated point estimates (posterior means) of population size and recombination parameters across test scenarios and embedding networks. As with CIs, mean relative errors use geometric means for population size parameters and the simple posterior mean for the recombination rate. Across scenarios the three embedding networks are roughly comparable in their accuracy for population size parameters, with the RNN producing the most accuracy population size estimates for two of the three scenarios, and the CNN performing best in the case of the linear decline. For recombination rate we observe similar variation, where there is no clear universally best embedding network; although for the linear growth scenario, the CNN embedding yields a considerably better recombination rate estimate than do the other networks.

#### Priors for Historical Population Size Inference

Prior specification plays a crucial role in Bayesian inference. To explore the impact of prior choice, we introduced a second prior for training the neural posterior estimator. As an alternative to the uniform prior, where each historical population size is sampled independently, we used a structured dependent prior. This alternative prior is designed to generate more contiguous and realistic population size histories by inducing auto-correlation between population sizes in adjacent time intervals; i.e., in the dependent prior, each population size depends on its preceding value, such that changes in population size between neighboring time windows are constrained to be at most a factor of 10 in either direction.

Both the independent and dependent priors were used to separately simulate training data corresponding to 21 population size parameters from log-spaced time windows extending to 130,000 generations ago. As input for the NPE we used, instead of features extracted by an embedding net, the site frequency spectrum and linkage disequilibrium summary statistics calculated for each simulated dataset. We then analyzed the performance of neural posterior estimators trained on the summary statistics generated by the uniform prior compared to the dependent prior by inferring the effective population size histories under six exemplary demographic scenarios. Regardless of the prior on the population size history, we use a uniform [10^−9^, 10^−8^] prior on the recombination rate.

Figure 9 illustrates the reconstructed effective population size histories for these scenarios. Both neural posterior estimators successfully capture the general demographic trends, with true population sizes largely falling within the 50% credible intervals. However, the choice of prior had a substantial impact on the inference quality and certainty.

**Figure 9:**
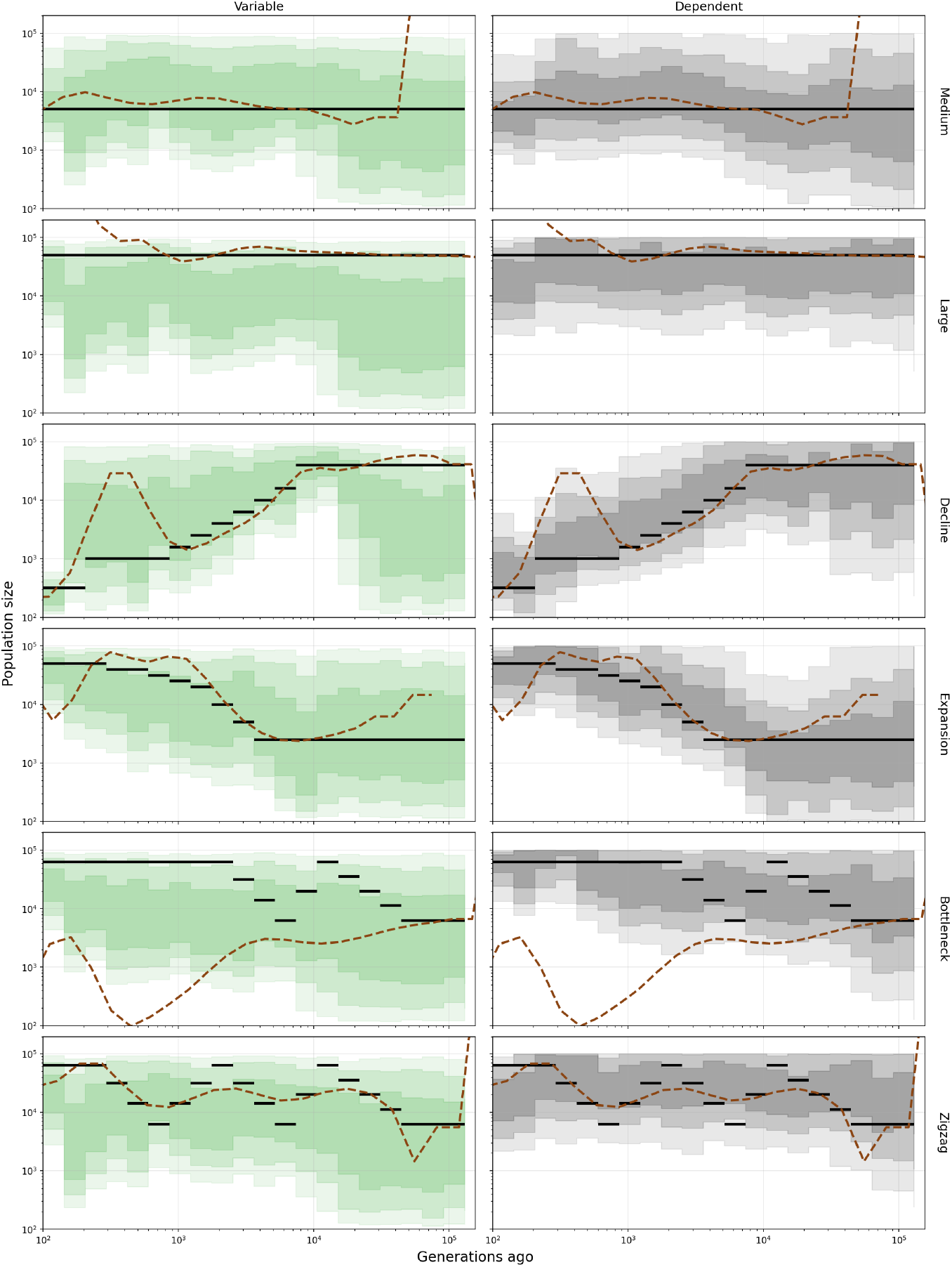
Inference of effective population sizes using variable and dependent prior under different scenarios. Scenarios shown in rows are: “Medium” and “Large” constant population sizes. “Decline” and “Expansion” where the population sizes slowly drop and increase. “Bottleneck” and “Zigzag” where population sizes fluctuate over time. MSMC2 estimates are shown as dashed lines for comparison.

In the “large” scenario, the neural posterior estimator trained on the dependent prior achieved a higher accuracy, when the one using uniform prior simply “missed” the ground truth in a large part of all time windows. We hypothesize that the uniform prior struggles to generate enough demographies with constant large population sizes, as every single historical population size window is drawn independently and uniformly, setting the population size more frequently to intermediate and low values. Therefore, scenarios close to the “large” scenario are less likely generated by the independent prior training data. In contrast, such scenarios occur more likely in the training data generated from the dependent prior, which improves the performance of NPE for these scenarios. In addition, the dependent prior also shows a relatively smooth and confident reconstruction of the “decline” and “expansion” scenarios.

Just as any other population genetic feature, the summary statistics used here provide less information for more ancient times (particularly beyond the MRCA), and consequently the neural posterior estimator trained on the log-uniform prior quickly reverts to the prior distribution in the ancient time windows. Meanwhile, the contiguous structure of the dependent prior leads the neural posterior estimator to project more constrained estimates backward in time. For the complex “bottleneck” and “zigzag” scenarios, both priors resulted in NPEs with reduced accuracy and confidence, struggling to fully resolve the population size fluctuations.

Similarly to the previous example, we quantify these performance differences using the mean relative errors in Figure S11. Consistent with the qualitative results, the neural posterior estimator trained on the dependent prior demonstrates lower relative error for effective population sizes estimation across all scenarios except for “zigzag”. In contrast, for the estimation of the recombination rate, the neural posterior estimator trained on the uniform prior consistently achieves higher accuracy. This might be caused by the highly variable effective population size histories sampled under the uniform prior, where information from neighboring time windows provides less information such that the neural posterior estimator might be encouraged to more robustly disentangle the genomic signals of recombination from those of population size.

### *Drosophila melanogaster* out-of-Africa model

As an example application to empirical data, we used our NPE pipeline to fit a joint demographic model for samples of African and European *Drosophila melanogaster*. In particular, we analyzed inbred samples from France and Cameroon, collected as part of the DPGP2 project (Pool et al., 2012), and fitted a modification of the Li and Stephan (2006) model. This model was originally used to represent the out-of-Africa expansion of *Drosophila*, and incorporates divergence from an ancestral population, migration, and exponential growth in the European lineage. Our modified model has seven free parameters: the ancestral population size, the effective sizes of the African and European populations following the split, the contemporary European effective size, the divergence time between populations, and asymmetric migration rates between the European and African populations. As described in the Methods section, we used an RNN embedding network trained on 300 Kb non-overlapping windows across chromosome 2L (we focus on 2L purely out of convenience and as a proof of principle).

In Figure 10 (Table S13), we show averaged posterior estimates of the seven demographic model parameters, with marginal distributions shown on the diagonal, and joint distributions shown in off-diagonal plots. Generally, all parameters show relatively narrow credible intervals, and distributions of population size estimates are particularly sharp, with the one exception of the founding size of the French population. Our population size estimates are in good agreement with those of Li and Stephan (2006), although we have used a different chromosome arm (the initial study used chromosome X) and different population samples (i.e., Li and Stephan used flies from the Netherlands and Zimbabwe). We estimate the current French effective population size is approximately 1 × 10^6^ and our estimate of the effective population size in Cameroon is appears to be a bit larger than that. We don’t find evidence in our estimates of as strong a bottleneck in the founding of the French population as was found in Li and Stephan (2006)— our estimate (posterior median) is on the order of *N*_*e*_ ∼ 82, 000 which is considerably larger than the estimate of *N*_*e*_ = 2, 200 reported in Li and Stephan. However the 95% credibility interval overlaps with the confidence interval on founding European *N*_*e*_ presented in that paper, and indeed our uncertainty of that parameter is larger than all other parameters examined. Our estimate of the split time between populations is on the order of t_*div*_ ∼ 180, 000 generations ago, and is thus quite comparable to estimates from Li and Stephan, who estimated divergence time between Netherlands and Zimbabwe to be approximately 158,000 generations ago. Examination of the joint posterior distributions reveals little posterior correlation among the majority of parameters, with most joint posteriors showing generally uncorrelated, elliptical shapes. A notable exception is the joint posterior between the current French effective population size and the migration parameter which determines the rate of influx (forward-in-time) from Cameroon to France, which show some negative correlation (Pearson correlation of *r* = − 0.4, pooling posterior samples over windows). This makes sense from the perspective of diversity—higher levels of contemporary genetic variation in France could be the result of larger effective population size or greater migration from Africa.

**Figure 10:**
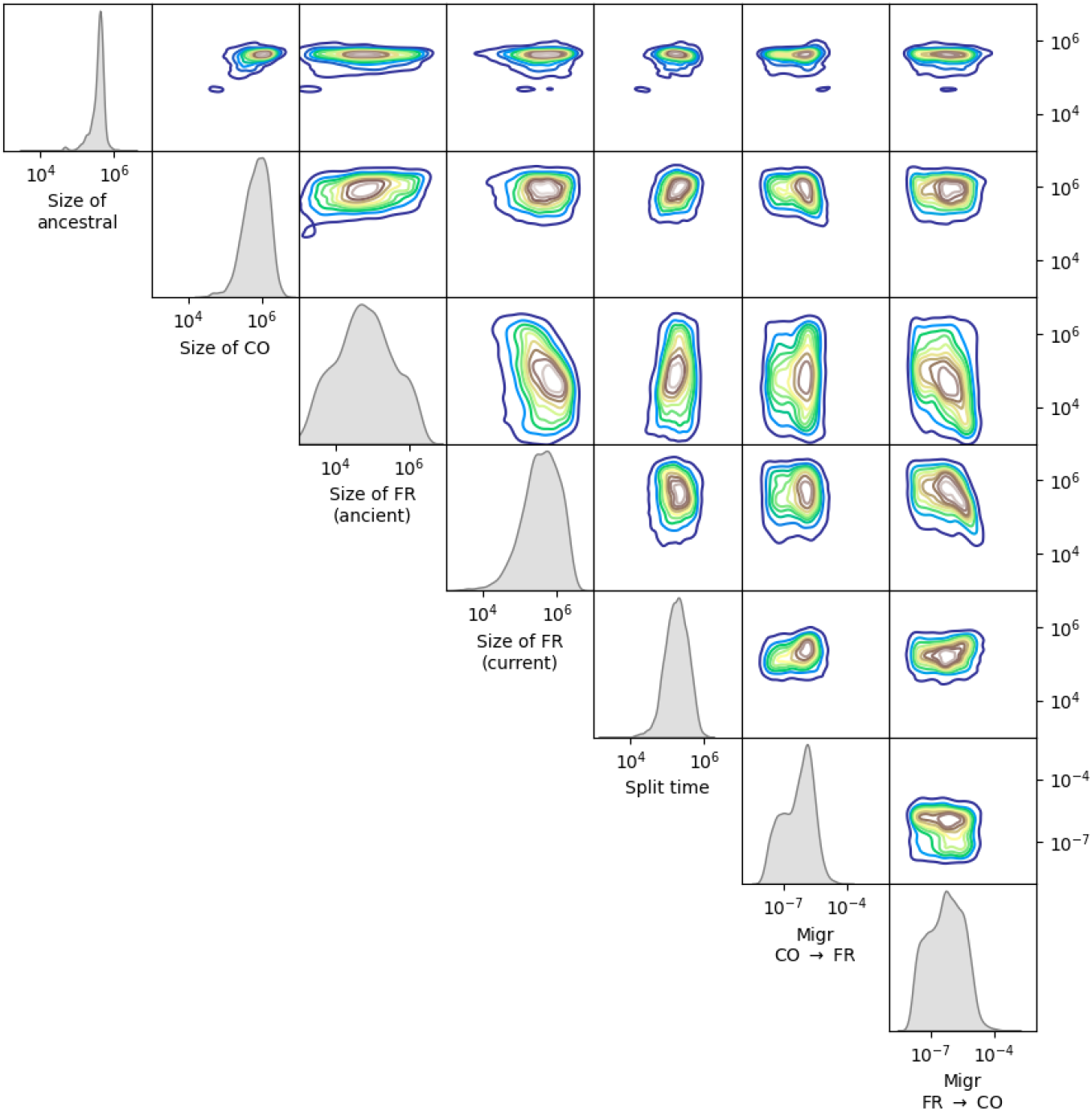
Joint posteriors for an *Drosophila melanogaster* out-of-Africa demographic model, fit via a neural posterior estimator with an RNN embedding network to 10 haploids from Africa (Cameroon) and 9 from Europe (France). Posteriors were estimated from unpolarised genotypes for 300Kb windows across 2L (see Fig S12, Fig 11), and then averaged to produce the contours shown here. Migration rates are backwards-in-time.

As we have estimated demographic parameters on a window-by-window basis, we can also examine how our estimates vary systematically across chromosome 2L (Fig S12). A few features are worth noting here. First, we see increased variation in effective population size parameters towards the distal (telomeric) and proximal (centromeric) regions of 2L. This is most likely due to the increased importance of linked selection in determining patterns of polymorphism in regions of low recombination, in the *Drosophila* genome (Aguade et al., 1989; Begun and Aquadro, 1992, e.g.,). We also see an interesting ‘blip’ in the estimates of split time and migration in the second-closest window to the centromere. We can only speculate at this point that this may be pointing to local genomic processes, such as an adaptive allele, but it perhaps echoes some of the unique dynamics found in centromere-proximal regions from humans (Langley et al., 2019, and C. Langley *pers. comm*. regarding flies).

To verify that the per-window variability in demographic parameter estimates is accurate, we performed posterior predictive simulations for each window, in which we drew samples from the per-window posterior, conducted coalescent simulations, and then summarized those simulations using a collection of summary statistics not used in estimation. The results of our posterior predictive checks are shown in Figure 11. As can be seen clearly, our posterior predictive simulations produce per-window variation in observed summary statistics that very closely matches the empirical summaries of the data. This gives us confidence that our NPE procedure is working well and providing meaningful fits to the data.

**Figure 11:**
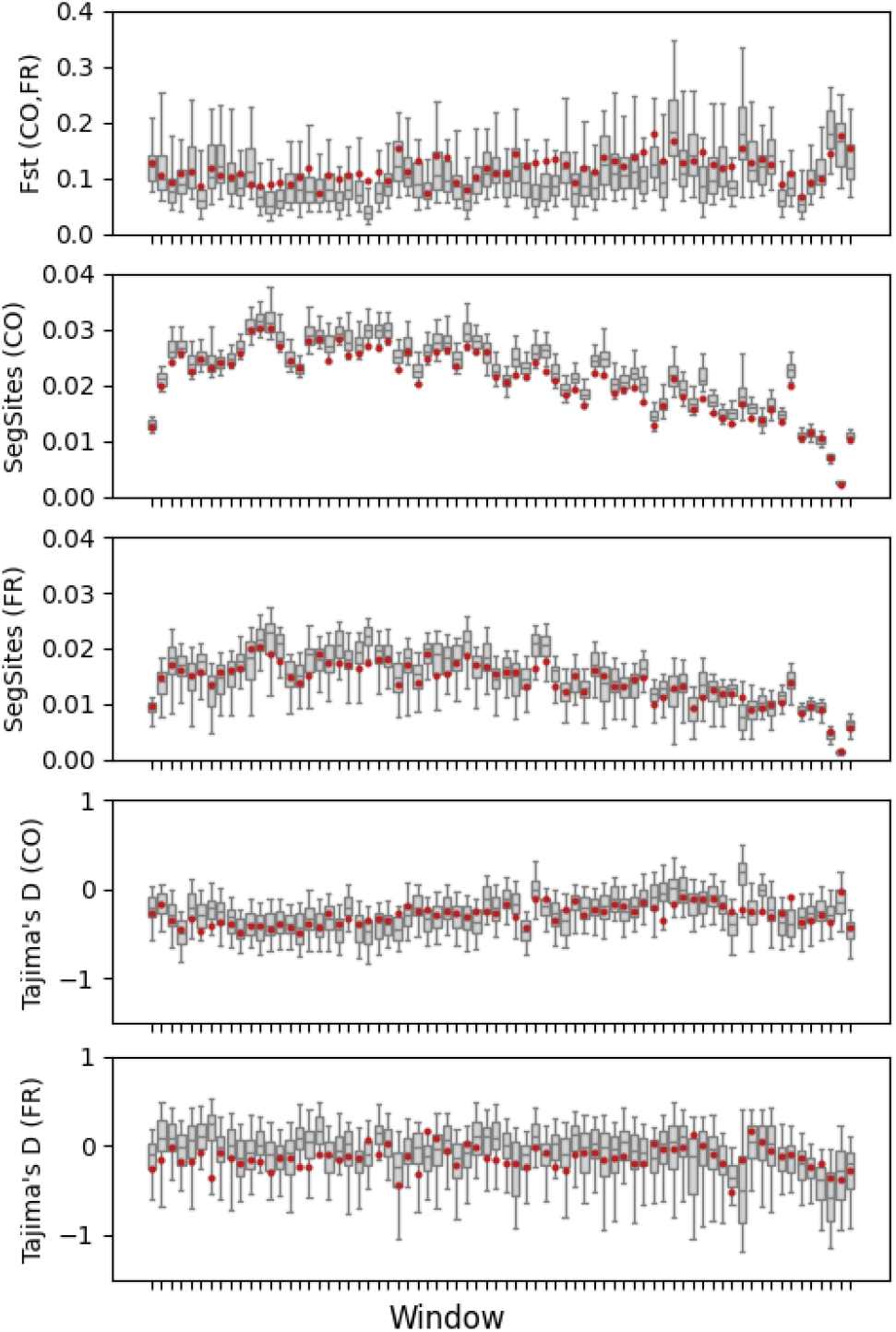
Posterior predictive checks for a *Drosophila melanogaster* out-of-Africa demographic model, fit via a neural posterior estimator with an RNN embedding network in 300Kb windows across 2L. For each window, demographic parameters were sampled from the posterior, genotype data was simulated with `msprime`, and posterior summary statistics (boxplots) plotted alongside to observed values (red points). The demographic parameter estimates in each window are shown in Fig S12.

## Discussion

In this study, we have demonstrated that neural posterior estimation (NPE) provides a powerful and flexible framework for population genetic inference that combines the principled uncertainty quantification of Bayesian methods with the computational efficiency and automatic feature extraction capabilities of deep learning. Our results across multiple inference tasks—recombination rate estimation, demographic parameter inference, and effective population size reconstruction—consistently show that NPE produces well-calibrated posterior distributions from both summary statistics and learned neural representations of raw genomic data.

The most striking advantage of NPE over existing methods is its ability to learn complex, non-linear posterior distributions directly from data. Unlike composite likelihood approaches such as `moments`, which rely on Gaussian approximations through asymptotic variance-covariance matrices, NPE captures the full shape of the posterior distribution. This was particularly evident in our bottleneck inference results (Figure 5), where the non-linear correlation between bottleneck timing (*T*) and intensity (*ν*) was accurately represented by NPE but led to miscalibrated confidence intervals when using Godambe information matrix approximations. This finding has important implications for demographic inference, where parameter correlations are common and often non-linear due to the complex relationship between demographic events and genetic patterns.

Compared to maximum likelihood based approaches, NPE offers dramatic computational savings through amortization. Once trained, an NPE model can generate posterior distributions for new observations almost instantaneously, whereas likelihood based-inference requires new optimization runs for each new obseration. Our recombination rate estimation results exemplify this advantage in a comparison to a previous method developed by our group (Adrion et al., 2020a): while parametric bootstrapping required 1,000 new simulations per genomic window to generate confidence intervals, the trained NPE model could produce equally well-calibrated credible intervals without any additional simulation. For genome-wide analyses involving thousands of windows, this represents a computational speedup of several orders of magnitude.

Furthermore, NPE’s flexibility in handling different data representations — from hand-crafted summary statistics to raw genomic data processed through embedding networks — allows researchers to choose the most appropriate representation for their specific problem. For instance, our results show that SFS summaries can be used effectively in conjunction with NPE, and thus can produce richer posterior representations than likelihood-based inference alone; however, incorporating linkage disequilibrium information through learned representations substantially improves inference accuracy and posterior concentration. This flexibility extends beyond current methods: as new network architectures are developed for genomic data, they can be readily incorporated into the NPE framework as embedding networks.

Our comparison of different data representations reveals important insights about information content in population genetic data. The superior performance of methods incorporating linkage information over SFS-alone approaches underscores that single-site summaries, while mathematically convenient, discard substantial information about population history. This finding aligns with recent theoretical work showing that linkage patterns contain independent information about demographic history, particularly for recent events and complex scenarios involving multiple populations (e.g., Jouganous et al., 2017).

The success of end-to-end learning approaches using CNNs and RNNs demonstrates that neural networks can automatically discover informative features from raw genetic data, potentially capturing patterns that would be difficult to encode in traditional summary statistics. However, our results also show that carefully chosen summary statistics can perform competitively with or potentially even outperform end-to-end approaches in some scenarios. Moreover, we can leverage theoretical results to improve the performance of machine learning methods based on predefined summary statistics (Burger et al., 2022) and interpret feature importance. This suggests that the optimal choice between summary statistics and end-to-end learning may be problem-specific, depending on factors such as the dimensionality of the parameter space, the complexity of the demographic model, existing theoretical insights, computational costs for summary statistics, and the amount of available training data.

### Challenges and Limitations

Despite these advantages, several challenges remain in applying NPE to population genetic problems. First, the quality of posterior estimates depends critically on the realism of the simulations used for training. While we used neutral coalescent simulations throughout this study, real genomic data is shaped by both demographic history and natural selection. Incorporating selection into the training simulations, while computationally more demanding, will be essential for applying NPE to empirical data where the effects of background selection and selective sweeps cannot be ignored. This challenge of model misspecification, of course, touches not just simulation-based methods of inference, as we present here, but also traditional, statistical methods which make assumptions about the distribution from which data is drawn.

Second, NPE requires choosing an appropriate prior distribution for the parameters of interest. While this is a general challenge in Bayesian inference, it is particularly acute for NPE because the method learns the posterior-to-prior ratio. Poorly chosen priors can lead to inefficient learning or biased posteriors (Figure S1). We primarily used uniform and log-uniform priors, which represents a relatively uninformative choice. However, we also showed that more complex and informative priors that incorporate biological knowledge can improve the performance (Figure 9). More broadly, the fidelity of the training simulations themselves is a limitation—for instance, our recombination rate examples use uniform rates across each simulated window, whereas real genomes exhibit substantial rate heterogeneity. When the generative process used for training diverges from the true data-generating process, the learned posteriors may not transfer faithfully to empirical applications.

Third, scaling to high-dimensional parameter spaces is a challenge shared by all inference approaches, though the bottlenecks differ. Likelihood-based methods such as `moments` become increasingly difficult to optimize as the number of parameters grows, because the likelihood surface develops ridges, saddle points, and near-flat directions that frustrate gradient-based search; moreover, the Gaussian approximation underlying information-matrix confidence intervals becomes less reliable when parameters interact in complex, non-linear ways. ABC suffers even more acutely: as the parameter dimension increases, the fraction of simulations falling near the observed data shrinks exponentially (the “curse of dimensionality”), so that prohibitively large simulation budgets are required to populate the posterior adequately (Prangle, 2015). NPE sidesteps the rejection-sampling bottleneck by learning a direct mapping from data to posterior, but it faces its own scaling challenge: the normalizing flow must represent an increasingly complex density, and training data requirements grow with dimensionality, creating a trade-off between the number of parameters one attempts to infer and the precision with which each can be estimated. In this study we successfully inferred up to 22 parameters (21 population sizes plus recombination rate), but many population genetic models of interest involve substantially more. Recent advances in normalizing flow architectures, sequential training schemes, and embedding-network design may help push this boundary, but further research is needed to determine the practical limits of NPE for complex models and to develop diagnostics that flag when posterior quality degrades in high dimensions.

### Implications for Population Genomics

The ability of NPE to provide rapid, well-calibrated uncertainty estimates has important implications for population genomic studies. In conservation genetics, where demographic inference informs management decisions, having reliable confidence intervals is crucial for risk assessment. NPE’s ability to quantify uncertainty while accounting for parameter correlations provides more reliable inference than methods assuming Gaussian posteriors.

For human population genomics, where complex models involving multiple populations, admixture events, and migration are the norm, NPE offers a path toward more realistic inference. The computational efficiency of amortized inference makes it feasible to analyze genomic data from thousands of individuals, potentially revealing fine-scale population structure and history that would be computationally prohibitive with ABC or composite likelihood methods.

Moreover, NPE’s modular structure—separating feature extraction (embedding networks) from density estimation (normalizing flows)—aligns well with the modular nature of population genetic models and simulators. Different embedding networks can be designed for different data types (SNPs, structural variants, microsatellites), while the same normalizing flow architecture can be used for posterior estimation. This modularity facilitates the development of specialized tools for different inference problems while maintaining a common statistical framework.

### Future Directions

Several promising directions emerge from this work. First, extending NPE to handle missing data and uncertainty in data preprocessing (such as genotype-calling errors) would increase its applicability to real datasets. Second, developing embedding networks specifically designed for population genetic data, perhaps incorporating known properties of the coalescent process, could improve both accuracy and interpretability.

The combination of NPE with model selection and model averaging procedures represents another important direction. While the technology for model selection through Bayes factors is well-established in the context of NPE, more sophisticated approaches could allow simultaneous inference of model structure and parameters (Schröder and Macke, 2024). This would be particularly valuable for scenarios where the demographic model itself is uncertain.

Finally, the integration of NPE with functional genomic data could enable joint inference of demographic history and selection, which is a particularly challenging and important goal (Boyko et al., 2008; Kim et al., 2017; Johri et al., 2020, 2021). By incorporating information about gene function, expression, and regulation into the embedding networks, it may be possible to disentangle the effects of demography and selection more effectively than current methods allow.

In conclusion, neural posterior estimation represents a significant methodological advance for population genetics, combining the best features of likelihood-based, ABC, and machine learning approaches. As genomic datasets continue to grow in size and complexity, methods like NPE that can efficiently extract information while providing principled uncertainty quantification will become increasingly important. The framework presented here provides a foundation for future developments in simulation-based inference, with the potential to enable more realistic and informative analysis of population genomic data.

## Acknowledgments

We thank Dan Schrider and Kent Holsinger for helpful comments on the manuscript. We thank Jakob Macke, Cornelius Schröder, and members of the Kern-Ralph colab for discussion and input on this project. We also thank Bruce Edelman who piloted this project as a programmer at UO. J.M., N.S.P, and A.D.K were supported in part by NIH grants R01HG010774, R01HG012473, and R35GM148253. F.B. was funded by the Deutsche Forschungsgemeinschaft (DFG, German Research Foundation) under Germany’s Excellence Strategy – EXC number 2064/1 – Project number 390727645, and EXC 2124 – Project number 390838134.

## Data Availability

All code for neural posterior estimation, training pipelines, and analysis scripts is available at https://github.com/kr-colab/popgen-npe. Simulation parameters and trained models sufficient to reproduce the analyses presented here are included in the repository.

