## Supplement for "Neural posterior estimation for population genetics"

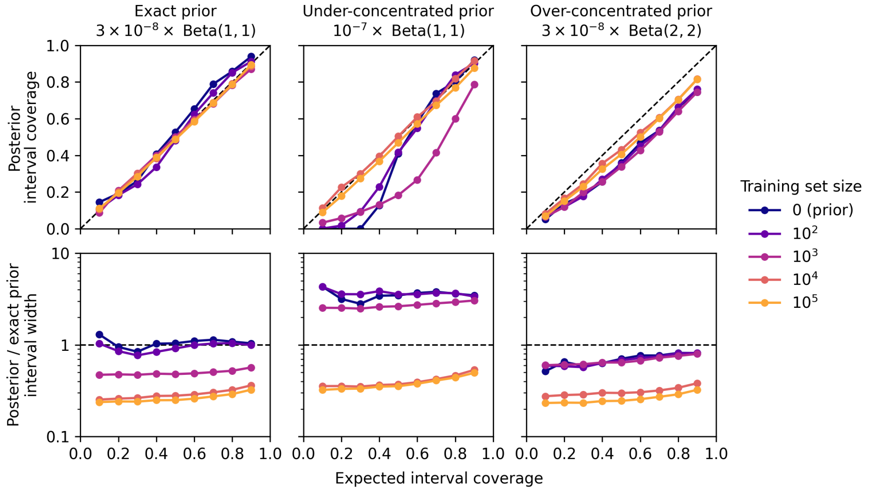


Figure S1: **Comparison of recombination rate posteriors across priors and training set sizes.** Here we show posterior coverage (top row) and concentration (posterior over prior interval width; bottom row) for NPE networks with an ReLERNN embedding layer when trained on different priors and numbers of training instances. All NPE were applied to the same test set with recombination rates simulated from a uniform distribution on $[0, 3 \times 10^{-8}]$. When trained on a prior matching the true generative process (left column), NPE posterior coverage matched expected coverage regardless of the number of training instances, and because concentrated relative to the prior by 1,000 training instances. In contrast, when trained on a uniform prior with wider support (middle column), NPE coverage and concentration were initially poor but because as well-calibrated as the "true NPE" by 10,000 training instances. Finally, when trained on an informative prior with the same support as the true generative process but a mode at $1.5 \times 10^{-8}$ (i.e. a Beta(2, 2) distribution rescaled to $[0, 3 \times 10^{-8}]$; right column), the NPE trained very quickly but produced posterior intervals that were too narrow on average.

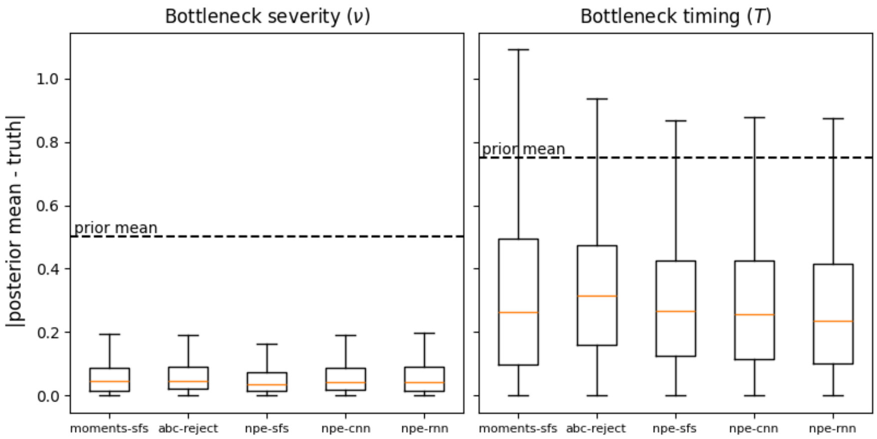


Figure S2: **Comparison of NPE point estimates to `moments` point estimates.** Here we show boxplot comparisons of NPE-derived point estimates to the `moments` point estimates across parameter space for $\nu$ and $T$ (left and right panels, respectively). Shown are the absolute error between the estimator and the truth across a validation set of 1000 simulations covering the prior parameter space. The dashed line is the average error when the prior mean is used as an estimator, representing a "worst-case" baseline.

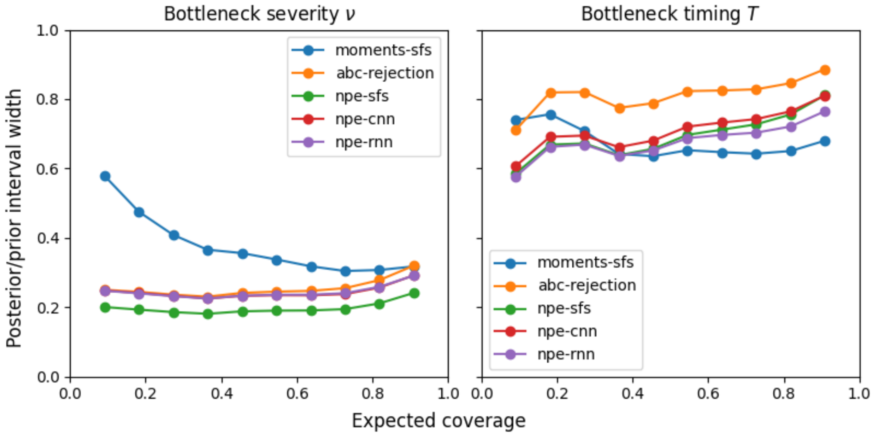


Figure S3: **Comparison of NPE posterior concentration to prior.** Here we show comparisons of the prior and posterior distributions for $\nu$ and $T$ for each of the three embedding approaches. We take the posterior interval width divided by the prior interval width as a measure of posterior "concentration", that is the gain in information from the data relative to the prior, across a range of interval widths. These are averaged across a validation set of 1000 simulations covering the prior parameter space. Additionally we show the interval ratio for the `moments` Godambe Information Matrix-corrected confidence intervals, where the prior in this case represents the bounds on the optimization.

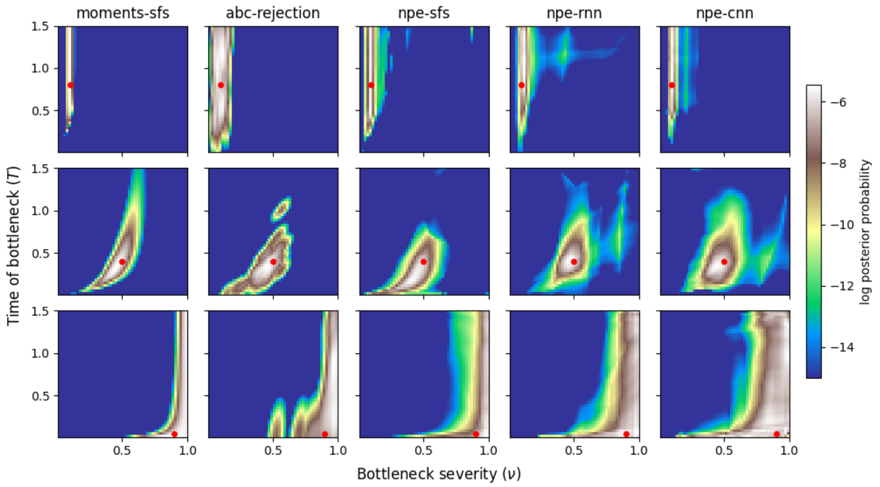


Figure S4: **Comparison of NPE posterior surfaces to `moments` likelihood surface (1 Mb).** Here we show the same comparison as in Figure 5 of the main text, but with 10x the sequence length (1Mb).

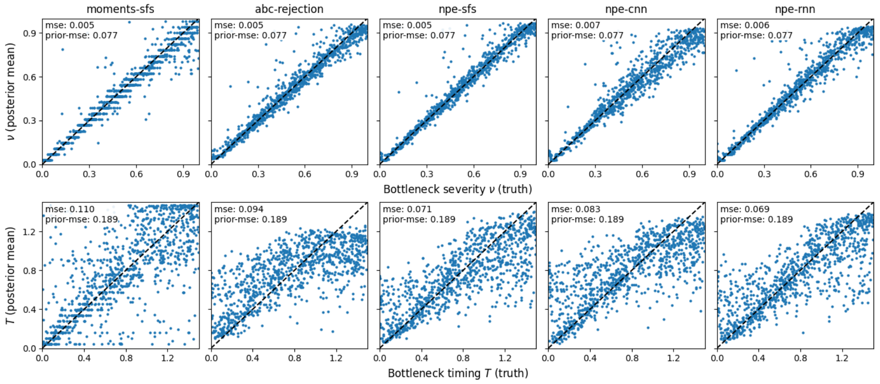


Figure S5: **Comparison of NPE point estimates to `moments` and ABC point estimates (1 Mb).** Here we show the same comparison as in Figure 6 of the main text, but with 10x the sequence length (1Mb).

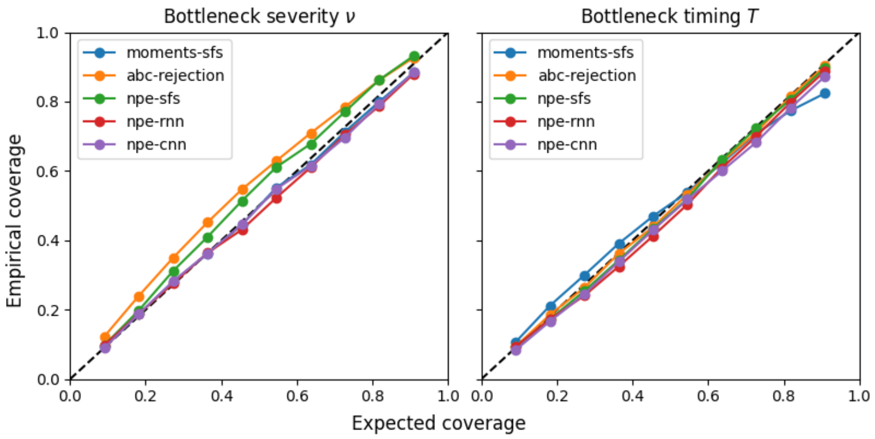


Figure S6: **Comparison of NPE posterior coverage (1 Mb).** Here we show the same comparison as in Figure 7 of the main text, but with 10x the sequence length (1Mb).

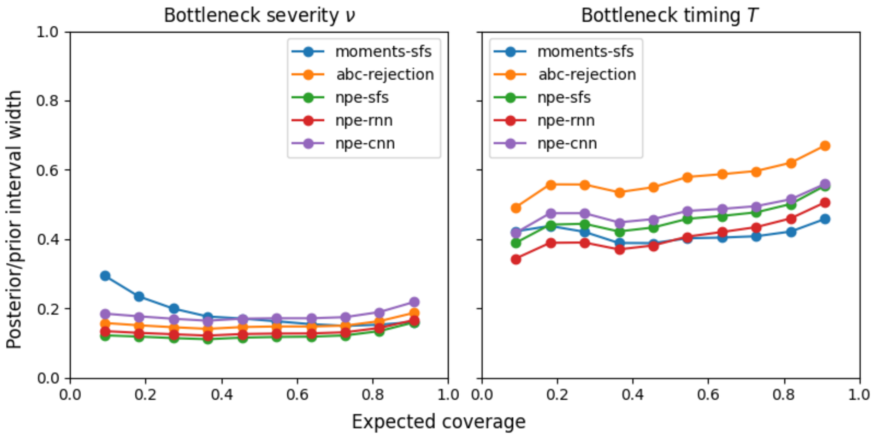


Figure S7: **Comparison of NPE posterior concentration to prior (1 Mb).** Here we show the same comparison as in Figure S3 of the main text, but with 10x the sequence length (1Mb).

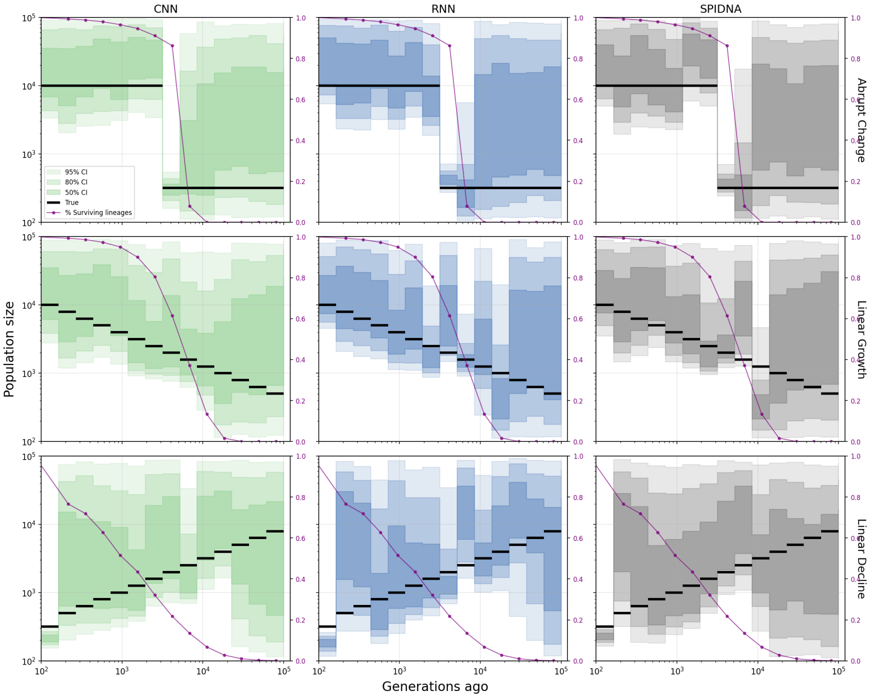


Figure S8: Benchmark of the neural posterior estimator on the task of inferring the effective population size over time using three different embedding networks. The three scenarios shown on rows are: 1) an abrupt change in population size, 2) a scenario of linear population growth, and 3) a scenario of exponential population growth. The three embedding networks shown on columns are: a CNN embedding network, a RNN embedding network, and the SPIDNA embedding network from (Sanchez et al., 2020). For each plot we show the true population size history (black line) and the inferred credible intervals (shaded regions) for the 15 time epochs. The purple line in each figure represents the percentage of remaining lineages at each time point within the test simulation.

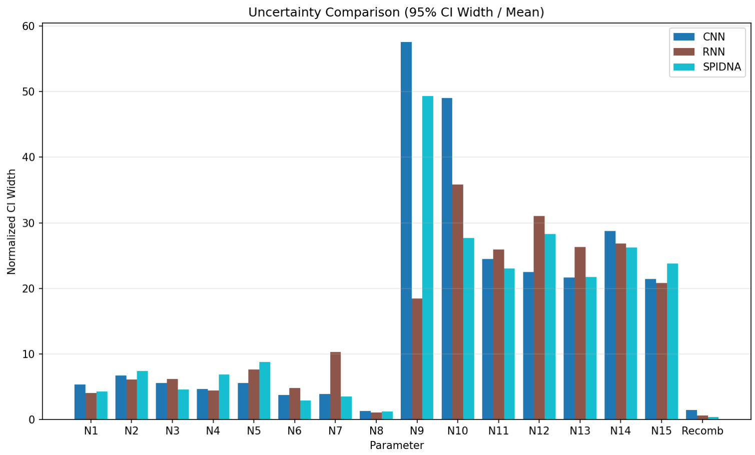


(a) Abrupt change

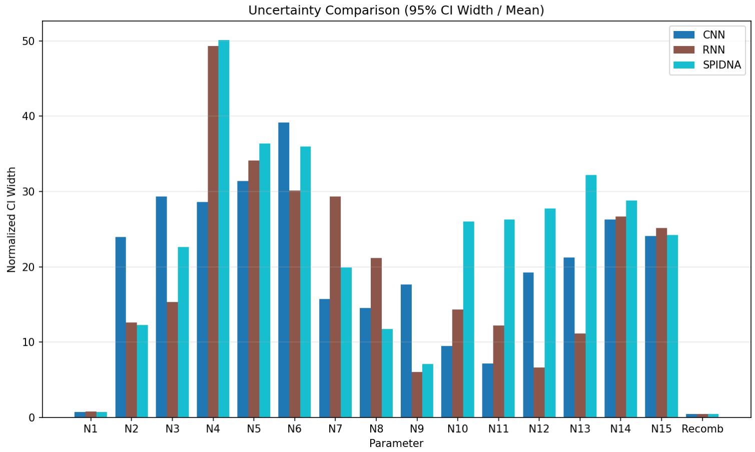


(b) Linear growth

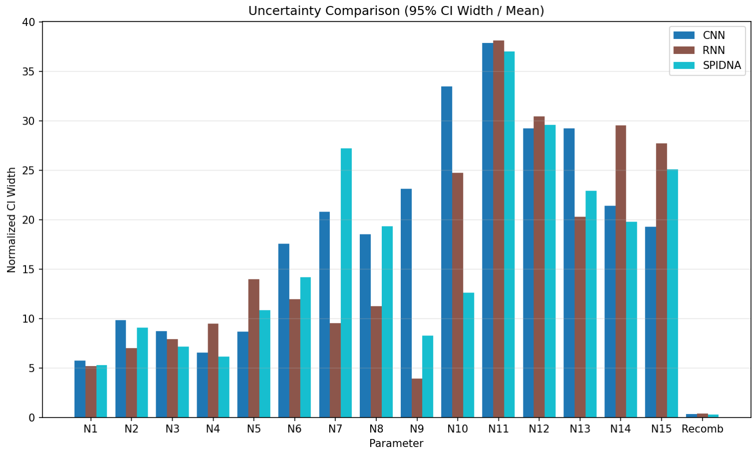


(c) Linear decline

Figure S9: Comparison of credible interval width across the three variable population size scenarios for each estimated parameter. Normalized credible interval width was calculated as the 95% credible interval width divided by the geometric mean for population size parameters and by the posterior mean for the recombination rate parameter.

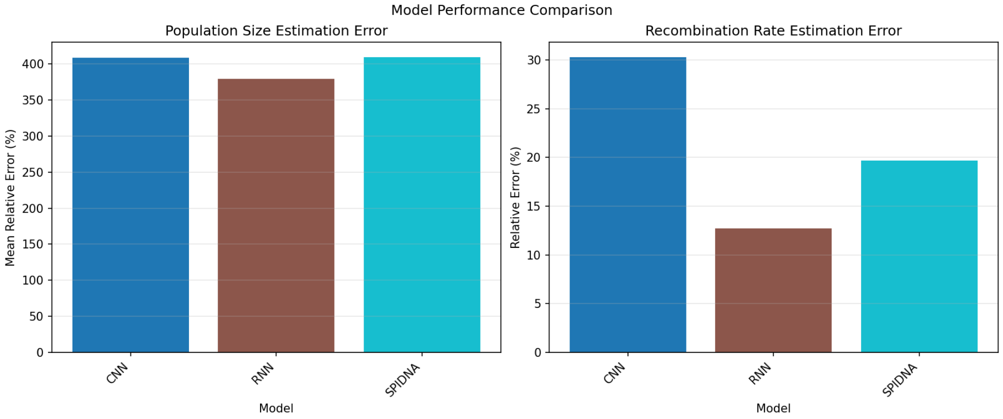


(a) Abrupt change

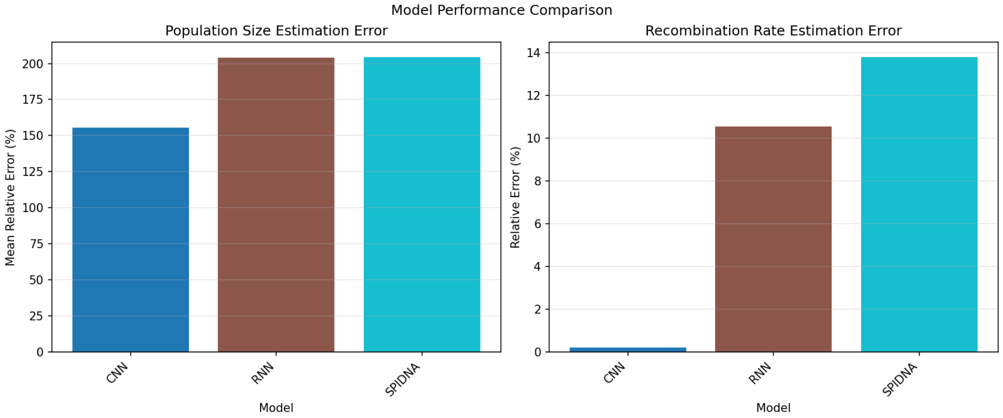


(b) Linear growth

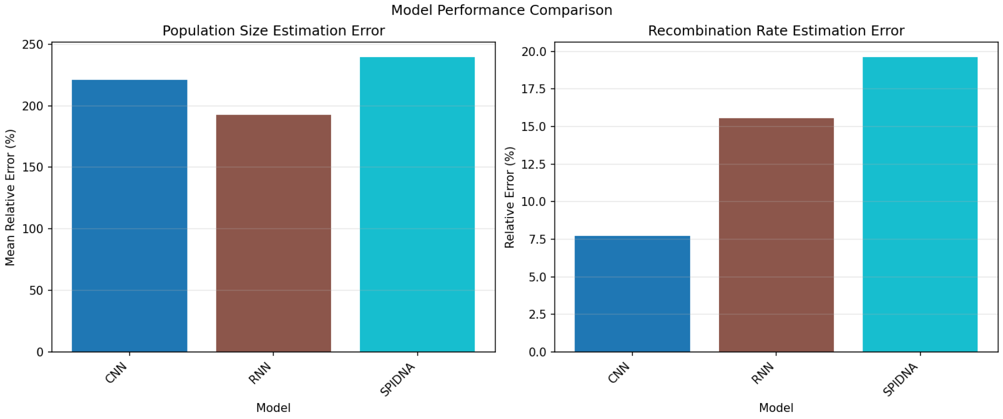


(c) Linear decline

Figure S10: Comparison of error in point estimation across the three variable population size scenarios. Left panel shows aggregated error across population size parameters, the right panel shows error for the recombination rate

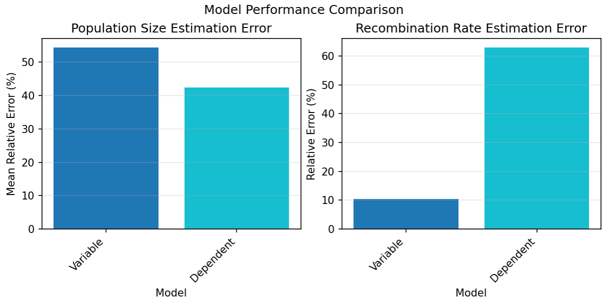


(a) Medium

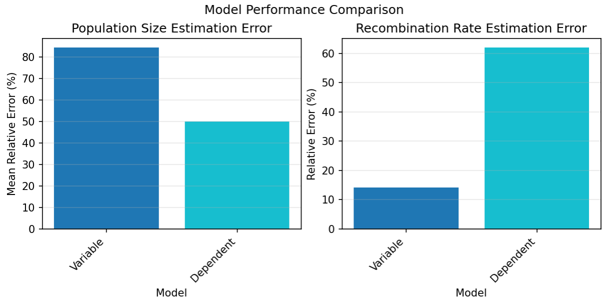


(b) Large

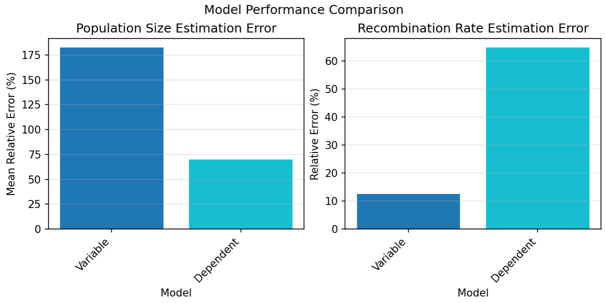


(c) Decline

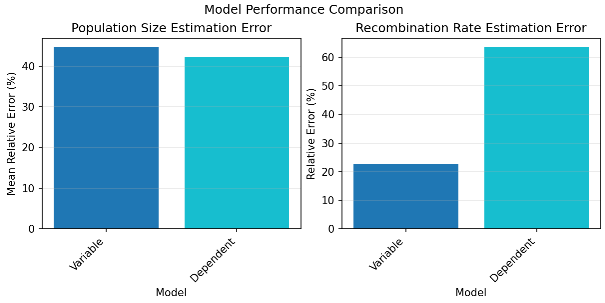


(d) Expansion

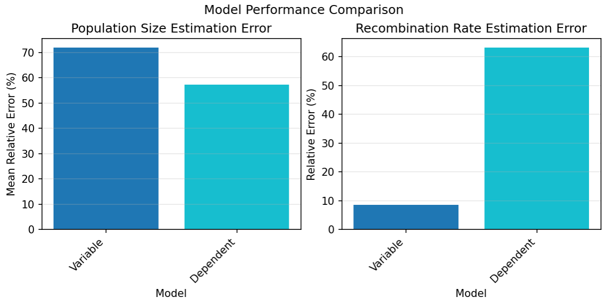


(e) Bottleneck

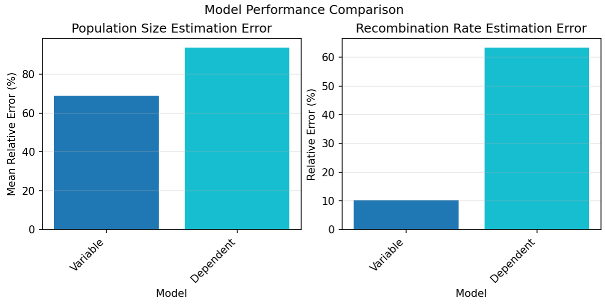


(f) Zigzag

Figure S11: Comparison of error in point estimation across six population size scenarios. Left panel shows aggregated error across population size parameters, the right panel shows error for the recombination rate

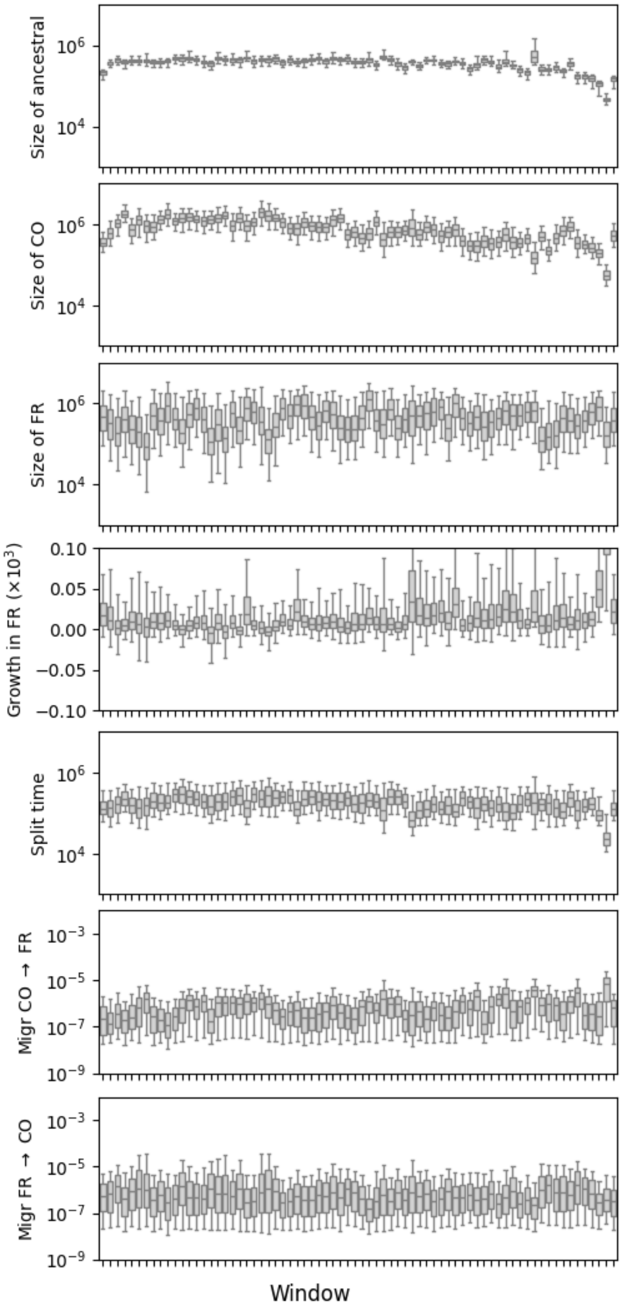


Figure S12: Marginal posterior distributions of parameters for a *Drosophila melanogaster* out-of-Africa demographic model, fit via a neural posterior estimator with an RNN embedding network in 300Kb windows across 2L. Migration rates are backwards-in-time.

| | | | Quantiles | | | |
|---|---|---|---|---|---|---|
| | Mean | Median | 0.025 | 0.25 | 0.75 | 0.975 |
| Size of ancestral | 380,867 | 389,288 | 109,605 | 305,789 | 451,407 | 632,108 |
| Size of CO | 888,294 | 740,976 | 125,023 | 416,628 | 1,218,365 | 2,420,950 |
| Size of FR (ancient) | 229,219 | 59,621 | 2,242 | 16,218 | 211,696 | 1,589,905 |
| Size of FR (current) | 619,429 | 395,504 | 27,203 | 172,051 | 863,027 | 2,283,723 |
| Split time | 216,028 | 176,029 | 40,075 | 107,993 | 285,626 | 593,438 |
| Migr CO $\rightarrow$ FR | 1.35e-06 | 6.04e-07 | 1.77e-08 | 1.17e-07 | 1.60e-06 | 6.81e-06 |
| Migr FR $\rightarrow$ CO | 2.56e-06 | 5.16e-07 | 1.67e-08 | 1.02e-07 | 2.11e-06 | 1.37e-05 |

Figure S13: Posterior summaries for parameters in the *Drosophila melanogaster* out-of-Africa demographic model, fit via a RNN embedding network.
